# A telomerase with novel non-canonical roles: TERT controls cellular aggregation and tissue size in *Dictyostelium*

**DOI:** 10.1101/556977

**Authors:** Nasna Nassir, Geoffrey J. Hyde, Ramamurthy Baskar

**Affiliations:** Department of Biotechnology, Bhupat and Jyoti Mehta School of Biosciences, Indian Institute of Technology-Madras, Chennai, India; Independent Researcher, 14 Randwick St, Randwick, New South Wales, Australia

## Abstract

Telomerase, particularly its main subunit, the reverse transcriptase, TERT, prevents DNA erosion during eukaryotic chromosomal replication, but also has poorly understood non-canonical functions. Here, in the model social amoeba *Dictyostelium discoideum,* we show that the protein encoded by *tert* has telomerase-like motifs and regulates, non-canonically, important developmental processes. Expression levels of wild-type (WT) *tert* were biphasic, peaking at 8 and 12 h post-starvation, aligning with developmental events, such as the initiation of streaming (∼7 h) and mound formation (∼10 h). In *tert* KO mutants, however, aggregation was delayed until 16 h. Large, irregular streams formed, then broke up, leading to small mounds. The mound-size defect was not induced when a KO mutant of *countin* (a master size-regulating gene) was treated with TERT inhibitors but anti-countin antibodies did rescue size in the *tert* KO. Further, conditioned medium from *countin* mutants failed to rescue size in the *tert* KO, but the converse experiment worked. These and additional observations indicate that TERT acts upstream of *smlA/countin* to regulate tissue size: (i) the observed expression levels of *smlA* and *countin,* being respectively lower and higher (than WT) in the *tert* KO; (ii) the levels of known size-regulation intermediates, glucose (low) and adenosine (high), in the *tert* mutant, and the size defect’s rescue by supplementing glucose or the adenosine-inhibitor, caffeine; (iii) the induction of the size defect in the WT by *tert* KO conditioned medium and TERT inhibitors. The *tert* KO’s other defects (delayed aggregation, irregular streaming) were associated with changes to cAMP-regulated processes (e.g. chemotaxis, cAMP pulsing) and their regulatory factors (e.g. cAMP; *acaA, carA* expression). Overexpression of WT *tert* in the *tert* KO rescued these defects (and size), and restored a single cAMP signalling centre. Our results indicate that TERT acts in novel, non-canonical and upstream ways, regulating key developmental events in *Dictyostelium*.

**Author summary:** When cells divide, their chromosomes are prone to shrinkage. This risk is reduced by an enzyme that repairs protective caps on each chromosome after cell division. This enzyme, telomerase, also has several other important but unrelated roles in human health. Most importantly, via one or other of its functions, both high and low levels of the enzyme can contribute to cancer. We have studied, for the first time, the roles played by telomerase in the life-cycle of the cellular slime mould, *Dictyostelium discoideum*, a model system with a rich history of helping us understand human biology. While we did not find any evidence of telomerase having the features typically needed to repair a chromosome, telomerase was necessary for many aspects of development. In forming the fruiting bodies that help *Dictyostelium* reproduce, a mutant that lacks telomerase miscalculates how big those bodies should be, and they end up being too small. Also, earlier, during an earlier stage, aggregation, the migration of cells that form each fruiting body is delayed and irregular. These results are significant because they show, for the first time, that a telomerase can influence cell migration and tissue size regulation, two processes involved in a wide range of cancers.

## Introduction

Each time a chromosome replicates, it loses some DNA from each of its ends. This is not necessarily problematic, because the chromosome is initially capped at each end by a sacrificial strand of non-coding DNA, a telomere [1–3]. Further instances of replication, however, can expose the coding DNA, unless the cell can keep repairing the shortened telomeres, by the action of the enzyme complex, telomerase. Accordingly, telomerase, whose main subunits comprise a reverse transcriptase (TERT), and the telomerase RNA component (TERC) [4], has much significance in the biology and pathology of multicellular organisms. As somatic tissues age, for example, telomerase is downregulated, and the resulting telomeric dysfunction can lead to chromosomal instability and various pathologies, including disrupted pregnancies and cancer [5–7]. In other cases, the upregulation of telomerase is also associated with, and a biomarker of, some cancers, because it allows the unchecked proliferation of immortalised tumour cells [6,8]. Telomerase also has many non-canonical roles, in which telomere maintenance, or even telomerase activity, is not required [9,10]. For example, telomerase is known to have non-canonical roles in neuronal differentiation [11], RNA silencing [12], enhanced mitochondrial function [13] and various cancers [9,14].

Our understanding of telomeres and telomerase began, and has continued to develop, through the study of model organisms such as *Drosophila, Zea mays, Tetrahymena*, yeast and mice [2,3,15–19]. One model system in which the possible roles of telomerase have not yet been addressed is *Dictyostelium discoideum*. This system has proved its usefulness in many contexts, including the study of human diseases [20–24]. One of its advantages is that the processes of cell division (i.e. growth) and development are uncoupled [25], making the organism a highly tractable system for the study, in particular, of differentiation and tissue size regulation [26–33]. In culture, when its bacterial food source is abundant, *D. discoideum* multiplies as single-celled amoebae. This leads to denser colonies, and exhaustion of the food supply. The rising concentration of a secreted glycoprotein, CMF, triggers the organism to switch to a multicellular mode of development [32,34]. With no resources for further cell proliferation, the amoebae move, in a radial pattern of streams, towards centres of aggregation. Rising levels of secreted proteins, of the counting factor (CF) complex [35,36], trigger a series of changes that lead to breaking up of the streams, which therefore no longer contribute cells to the original aggregate. Each aggregate, which will typically contain 20, 000 to 100,000 cells [37], now rounds up into a mound, which then proceeds through several life-cycle stages, finally forming a spore-dispersing fruiting body about 1-2mm high [32,38]. Mounds can also develop from the breaking-up of a large stream (or aggregate), a process similarly regulated by CF [27,39]. The generic term, ‘group’, can be used to address the fact that mounds develop from clusters that arise in these slightly different ways, but in this paper we will refer to ‘mounds’. Some of the processes and regulators involved in our very abbreviated account of the life-cycle are shown in Fig 1, which focuses on those elements examined in this study.

**Fig 1.**
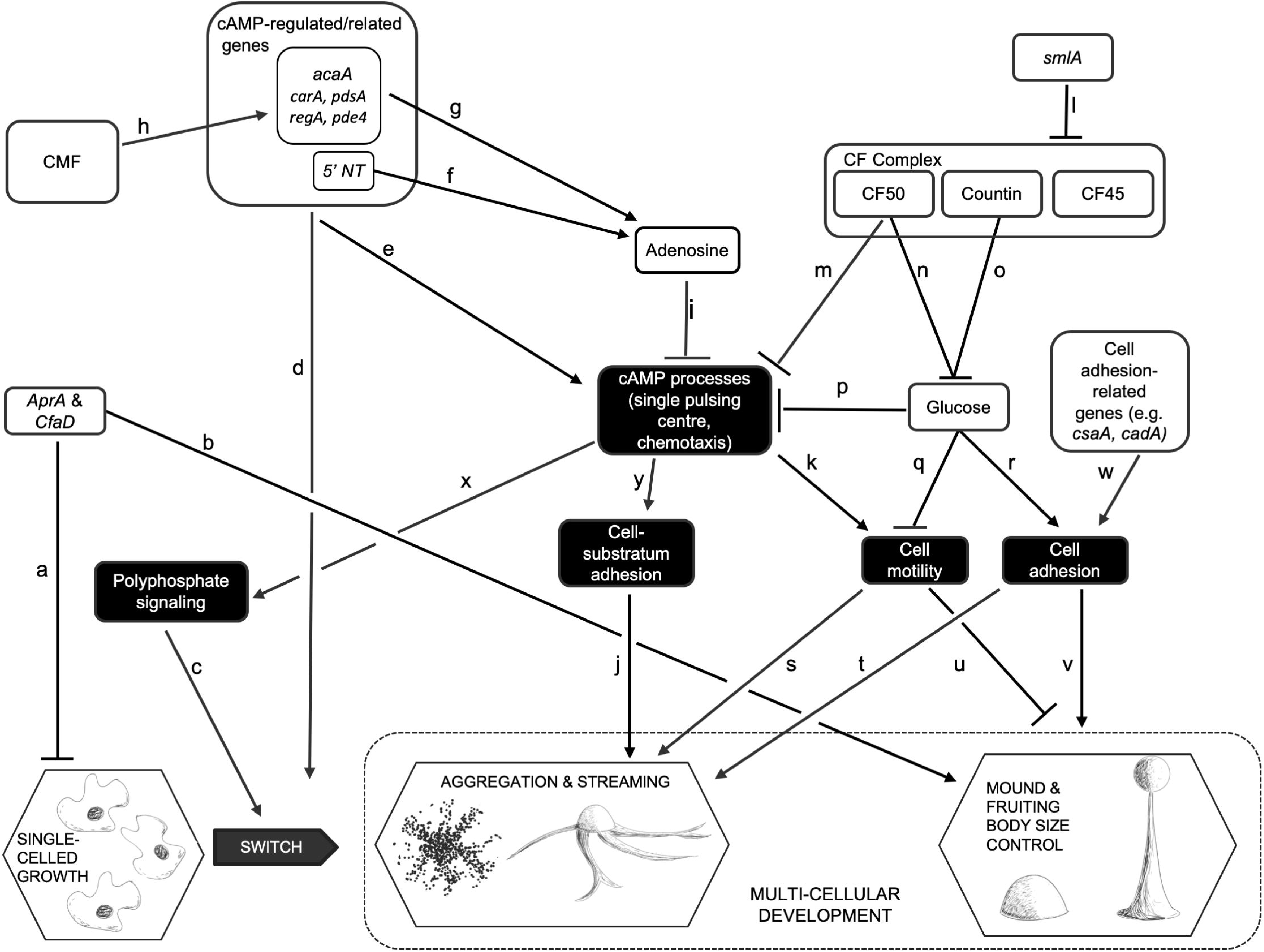
Some of the events, processes and regulators of growth and development in *D. discoideum.* This figure depicts only a small number of the hypothesized regulatory pathways of *Dictyostelium* growth and development, focusing on those that were examined experimentally in this study. A line ending in an arrowhead suggests that the first element directly or indirectly promotes the activity or levels of the second; inhibition is suggested by a line ending in a cross-bar. Published works that report on the nature of each pathway within the network are as follows: a[29], [40]; b[29]; c[66]; d[52], [53], [54]; e[61], [82], [83], [84]; f[85]; g [86], [87]; h[88–90]; i[91], [33]; j[77]; k[92], [93], [94]; l[26], [39]; m[27]; n[35]; o[95], [96]; p[67]; q[67], [75]; r[67]; s[97]; t[75]; u[65]; v[98], [39]; w[99], [100]; x[66]; y[101].

In addition to being uncoupled from growth, development in *D. discoideum* has other features that make it potentially useful as a model system for the understanding of telomerase-based pathologies, in particular cancers that arise from disruption of non-canonical functions. First, as indicated in Fig 1, development in *D. discoideum* depends on properly regulated cell motility and cell adhesion, two processes fundamental to metastasis. Second, the switch to multicellular development, and the control of aggregate, mound and hence fruiting body size are influenced by various secreted factors that, respectively, promote aggregation and regulate tissue size, in ways analogous to the regulation of tumour size by chalones [40,41]. Third, a putative TERT has been annotated in the *D. discoideum* genome. It is not known if the RNA component of telomerase (TERC) is present [42] and, in any case, extrachromosomal rDNA elements at the ends of each chromosome in *D. discoideum* suggest a novel telomere structure [43]. Thus, telomerase in this organism may have a separate mechanism for telomere addition or might have non-canonical roles. As yet, however, there have been no functional studies of TERT reported for *D. discoideum*.

In this study, we characterize the gene *tert* in *D. discoideum*, showing that it has both RT and RNA binding domains. We describe the pattern of *tert*’s expression levels during all stages of development, assay for any canonical telomerase function, and examine the effects of knocking out the gene’s function on development. The *tert* mutant exhibits a wide range of developmental defects, suggesting that wild-type TERT targets multiple elements of the regulatory network depicted in Fig 1. Most interestingly, these defects, and the results of experiments by which we attempt to rescue, or phenocopy, the *tert* KO phenotype, suggest that this telomerase influences the activity of *smlA*, and processes downstream of it. *Tert* thus emerges as the most upstream gene involved in the cell-counting pathway identified to date, and its overall influence indicates that, despite having no obvious canonical activity, a telomerase can nevertheless play major regulatory roles by virtue of its non-canonical targets.

## Results and discussion

### *D. discoideum* expresses *tert*, a gene encoding a protein with telomerase motifs

Extending previous predictions of *tert* encoding a protein with telomerase motifs [44], our use of the Simple Modular Architecture Research Tool (http://SMART.embl-heidelberg.de) and UniProt (Q54B44) revealed the presence of a highly conserved reverse transcriptase domain and a telomerase RNA binding domain (S1 Fig). These are characteristic of a telomerase reverse transcriptase [45], supporting the idea that the gene we characterize below indeed encodes for TERT. The protein shares 23% and 18.7% identity with human and yeast TERT protein respectively (Pairwise sequence Alignment-Emboss Needle).

Telomerase activity, if any, is ascertained by performing a Telomeric Repeat Amplification Protocol (TRAP) assay. However, while human cell lines (HeLa, HEK) showed telomerase activity, we did not detect any telomerase activity in *D. discoideum* cell extracts (S2 Fig). This concurs with previous findings, namely that the telomeres of *D. discoideum* have a novel structure [46], and that, in other organisms, TERT has several non-canonical roles [11–13].

### Constitutive expression of telomerase during growth and development in *D. discoideum*

In humans, telomerase expression is reported to be low in somatic cells compared to germline and tumour cells [47]. To ascertain if *tert* expression is differentially regulated during growth and/or development, we performed qRT-PCR using RNA from different developmental stages (0, 4, 8, 10, 12, 16 and 24 h after starvation). *Tert* expression is higher in development than during growth, (8h and 12 h) (Fig 2), implying that *tert* plays a prominent role beyond the point at which *D. discoideum* is responding to starvation. Expression also shows a marked biphasic pattern, with the first peak at 8h (when streams are forming), a big dip during stream breaking (10h) and then rising gradually again to peak at about the time of mound formation (16h).

**Fig 2.**
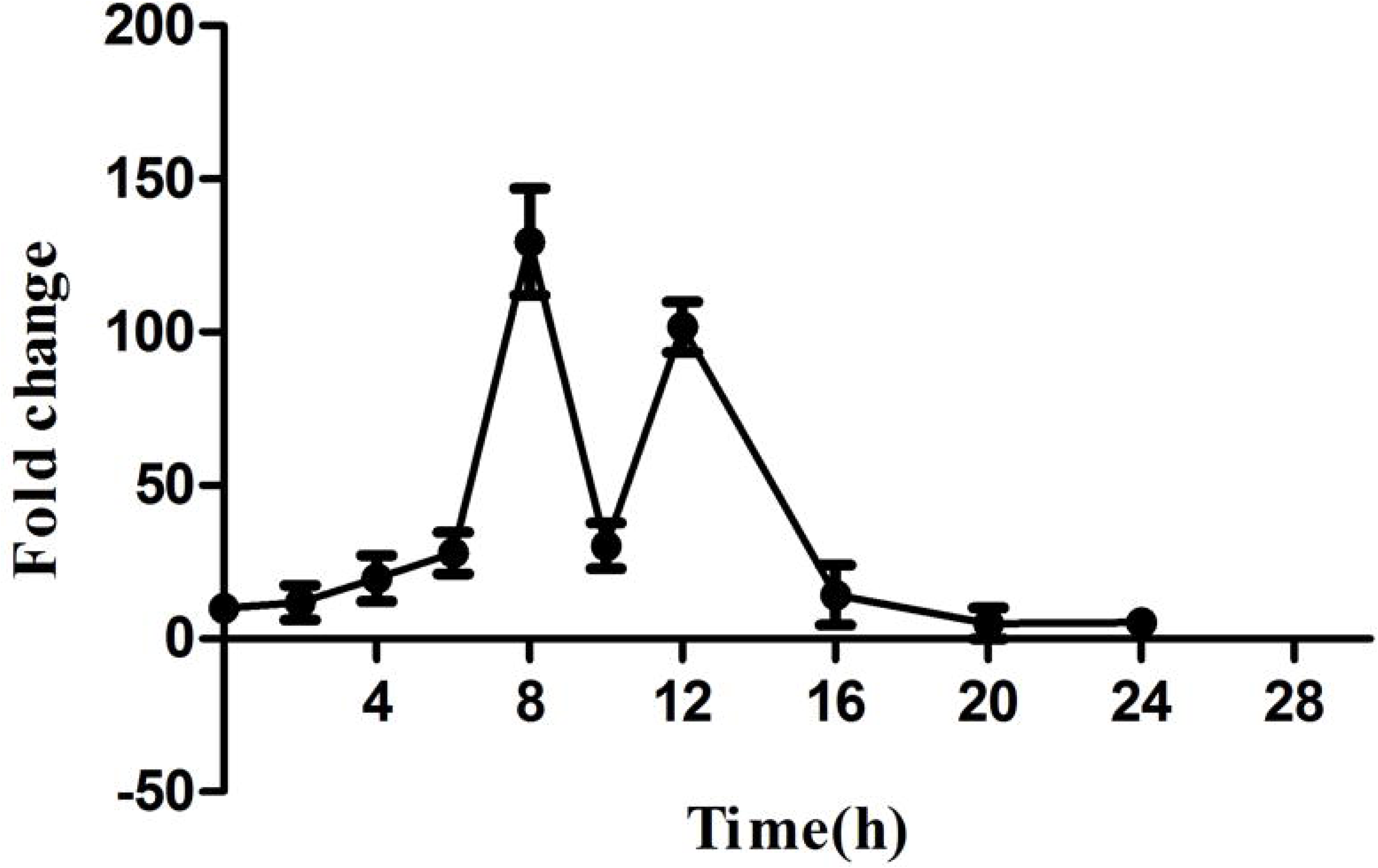
*Tert* expression during growth and development in *D. discoideum*. *Tert* is a single copy gene in *Dictyostelium.* Total RNA was extracted from *Dictyostelium* strain AX2 during vegetative growth and development. To analyze *tert* expression, qRT-PCR was carried out and fold change was calculated. *rnlA* was used as a control. Time points are shown in hours (bottom). qRT-PCR was carried out thrice. Error bars represent the mean and SEM.

### *tert* KO leads to delayed development, irregular streaming, and smaller mounds and fruiting bodies

To understand the possible non-canonical roles of *tert* in development in *D. discoideum, tert* KO cells generated by homologous recombination were seeded at a density of 5×10^5^ cells/cm^2^ on non-nutrient buffered agar plates and monitored throughout development. While aggregates appeared by 8 h in the wild-type, and streams began to break at 10 h, in the mutants there was a further 8 h delay before aggregates were seen, and stream breaking began at about 18 h. Wild-type cells formed long streams of polarized, elongated cells leading to aggregation but *tert* KO cells did not form well-defined streams, failing to aggregate even at 5×10^4^ cells/cm^2^ (wild-type cells aggregated at a density of 2×10^4^ cells/cm^2^), suggesting an inability to respond to aggregation-triggering conditions (S3 Fig). The mutant’s streams were also larger (Fig 3A). In contrast to streams moving continuously towards the aggregation centre in WT, *tert* KO streams break while they aggregate (Supplementary Videos 1 and 2). They did eventually form aggregates, largely by clumping. During the early stages of aggregate formation, the number of aggregates formed by the *tert* KO was only 10% of that formed by WT (Fig 3B, p<0.0001). Due to uneven fragmentation, the late aggregates were also of mixed sizes. The *tert* KO cells did eventually form all of the typical developmental structures, but by the mound stage, continued fragmentation had resulted in the mounds being more numerous, and smaller, on average, than in the WT. This was also the case for fruiting bodies. Thus, with reference to Fig 1, *tert* appears to play roles in multiple aspects of *Dictyostelium* development: the timing of aggregation; streaming, and the regulation of the size of the mound and fruiting body (Table 1).

**Table 1a.** Phenotypic differences between wild-type and *tert* KO development of *D. discoideum*, and some possible causal factors (Part 1 of 2)

|  | Timing of delay to aggregation | Streaming and Aggregation | Mound (and fruiting body) size | Mound-size regulation pathway |  |  |
| --- | --- | --- | --- | --- | --- | --- |
|  |  |  |  | *smlA* expression levels | *Countin* expression levels | Glucose levels |
| Wild-type cells | Normal | Normal | Normal | Normal | Normal | Normal |
| *tert* KO cells | Delayed (by 8h) | Fragmented, uneven aggregates | Small | Low | High* | Low* |

**Table 1b.** Phenotypic differences between wild-type and *tert* KO development of *D. discoideum*, and some possible causal factors (Part 2 of 2)

|  | cAMP-related factors (streaming & delay regulation) |  |  |  |  | Adhesion-related factors (in streaming and mound-size regulation) |  |  | Delay-specific regulation |
| --- | --- | --- | --- | --- | --- | --- | --- | --- | --- |
|  | Levels of cAMP-related genes: *acaA*, *carA*, *5'NT*, *pdsA*, *regA*, *pde4* | Adenosine/ 5'NT Levels | cAMP centres | cAMP levels | cAMP-based chemotaxis | *csaA*, *cadA* expression levels | Cell-cell adhesion | Cell-substratum adhesion | Poly-phosphate levels |
| Wild-type cells | Normal | Normal | One | Normal | Normal | Normal | Normal | Normal | Normal |
| *tert* KO cells | Low at 4h-10h; High by 12h | High/high during stream break; low after. * | Many* | Low* | Abnormal | Low | Low | Low | Low |

**Fig 3.**
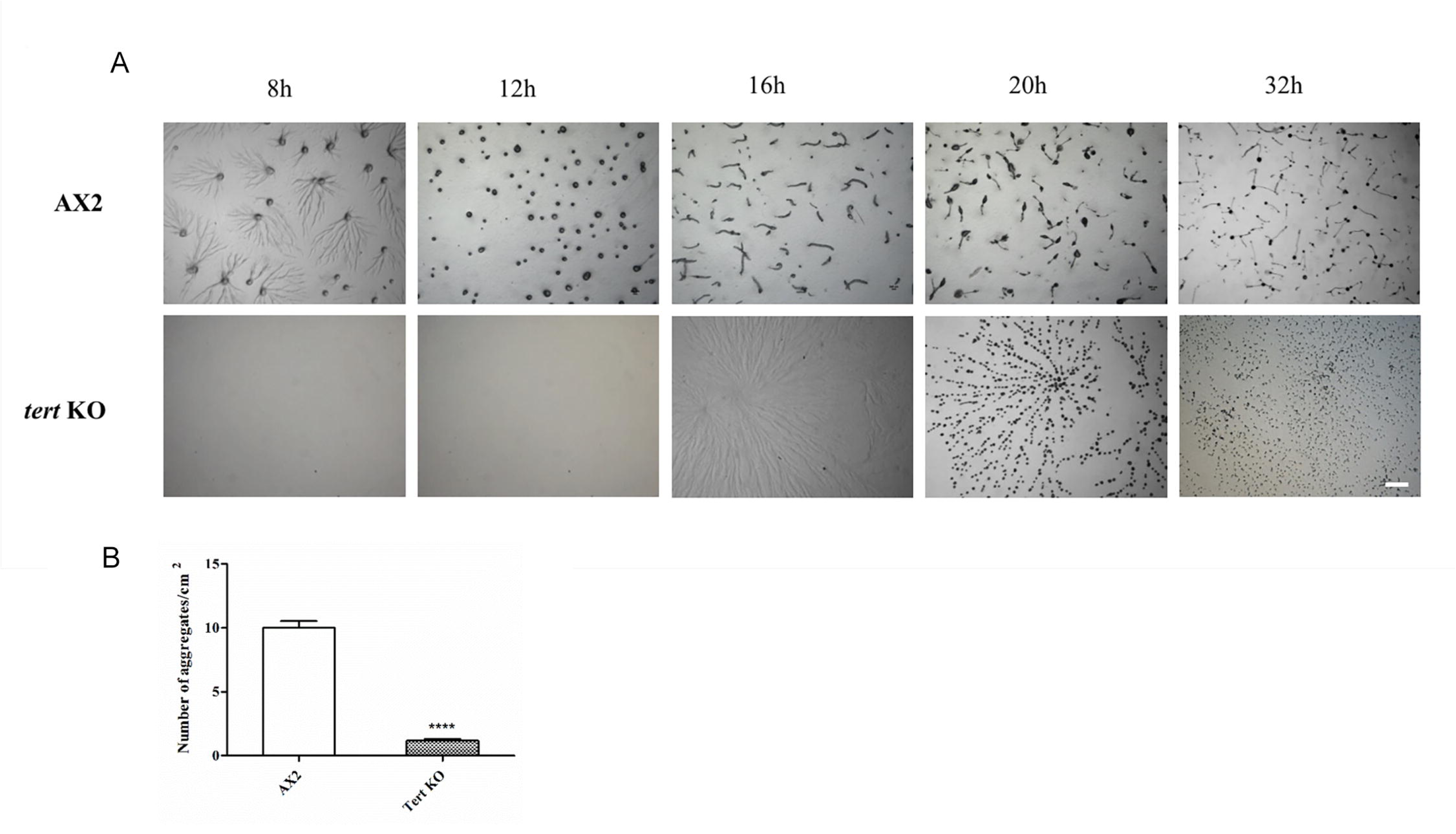
Developmental phenotype of *tert* KO. (A) AX2 and *tert* KO cells plated on 1% non-nutrient KK_2_ agar plates at a density of 5×10^5^ cells/cm^2^ were incubated in a dark, moist chamber. After 16 hours, large aggregate streams were formed in *tert* KO. The time points in hours are shown at the top. Scale bar-0.5 mm. (B) Quantitative measurement of aggregation. The number of aggregates was counted per centimetre square area. Level of significance is indicated as *p<0.05, **p<0.01, ***p<0.001, and ****p<0.0001.

### Many processes and regulators are potentially involved in the phenotypic changes of the *tert* KO

Given the wide-ranging phenotypic defects seen in the *tert* KO, it seemed likely that *tert* is a master regulator, affecting many of the processes and regulators depicted in Fig 1. We thus monitored the activity or levels of a number of those elements, comparing the wild-type and *tert* KO (summarised in Table 1). As that summary shows, the *tert* KO showed significant changes from the wild-type in three broad areas: components of the mound-size regulation pathway; cAMP-related processes/regulators; adhesion-related processes/regulators. As is clear from Fig 1, the factors that influence these features overlap considerably, both in terms of interacting with each other, and in regulating more than one of the various developmental stages disrupted in the *tert* KO. Nevertheless, we think it is useful to consider each of them in turn. As we do so below, we describe a series of experiments that largely fall into two broad categories, as shown in summary form in Tables 2 and 3: Those that attempt to rescue the normal phenotype in *tert* KO cells (Table 2); and those that attempt to phenocopy, or induce, the *tert* KO phenotype in wild-type cells (Table 3). First, however, we describe some experiments that support the direct involvement of *tert* in the effects already noted.

**Table 2.** Attempts to rescue normal phenotype (or aggravate^#^ the KO phenotype) in *D. discoideum tert* KO cells.

|  | Timing of delay to aggregation | Streaming and Aggregation | Mound and fruiting body size |
| --- | --- | --- | --- |
| Overexpression of WT *tert* | Rescue | Rescue | Rescue |
| Overexpression of human *tert* | No Rescue | No Rescue | No Rescue |
| WT cells (10-50% of total cells) | Full rescue at 50% | No Rescue | No Rescue |
| WT Conditioned Medium | No Rescue | No Rescue | No Rescue |
| Anti-countin or anti-CF50 antibodies | No Rescue | Part Rescue | Part Rescue |
| Anti-CF45 antibodies | No Rescue | No Rescue | No Rescue |
| *Countin* KO Conditioned Medium | No Rescue | No Rescue | No Rescue |
| 1mM glucose | No Rescue | Rescue | Rescue |
| Caffeine | No Rescue | Rescue | Rescue |
| cAMP pulsing | No Rescue | No Rescue | No Rescue |
| 8-Br-cAMP | No Rescue | No Rescue | No Rescue |
| Anti-AprA, anti-CfaD antibodies | No Rescue | No Rescue | No Rescue |
| *tert* KO Conditioned Medium # | No aggravation<br>(i.e. 8h delay typical of *tert* KO) | Aggravation | Aggravation |

Green shading indicates full or partial rescue of normal (wild-type) levels or activity by a treatment applied to *tert* KO cells. Red shading indicates no rescue. The final row refers to an attempt to exacerbate the KO phenocopy.

**Table 3.**
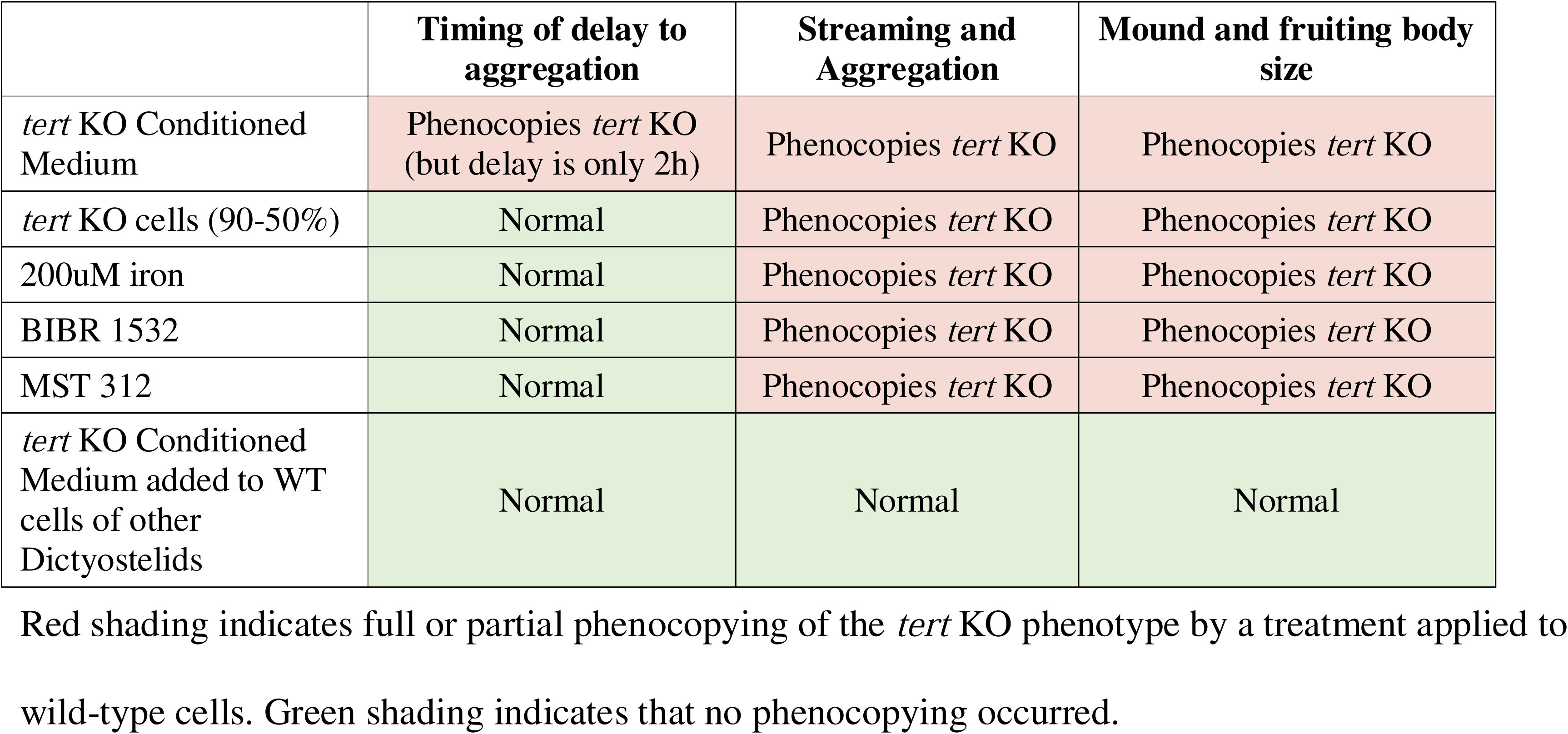
Attempts to phenocopy the *tert* KO phenotype in wild-type *Dictyostelium* cells.

|  | Timing of delay to aggregation | Streaming and Aggregation | Mound and fruiting body size |
| --- | --- | --- | --- |
| *tert* KO Conditioned Medium | Phenocopies *tert* KO (but delay is only 2h) | Phenocopies *tert* KO | Phenocopies *tert* KO |
| *tert* KO cells (90-50%) | Normal | Phenocopies *tert* KO | Phenocopies *tert* KO |
| 200uM iron | Normal | Phenocopies *tert* KO | Phenocopies *tert* KO |
| BIBR 1532 | Normal | Phenocopies *tert* KO | Phenocopies *tert* KO |
| MST 312 | Normal | Phenocopies *tert* KO | Phenocopies *tert* KO |
| *tert* KO Conditioned Medium added to WT cells of other Dictyostelids | Normal | Normal | Normal |

Red shading indicates full or partial phenocopying of the *tert* KO phenotype by a treatment applied to wild-type cells. Green shading indicates that no phenocopying occurred.

### Support for the involvement of *tert* itself in the *tert* KO

To support the idea that the changes observed in the *tert* KO are, in the first instance, due to changes involving *tert* itself, and not some other factor, we took two approaches: Overexpression of *tert,* and the use of TERT inhibitors. Most importantly, overexpression of wild-type TERT (act15/gfp::*tert*) in *tert* KO cells rescued all three of the phenotypic defects (Fig 4A, Supplementary Video 3; Table 2), suggesting that the *tert* KO phenotype is not due to any other mutation. Next, we treated wild-type cells with structurally unrelated TERT specific inhibitors, BIBR 1532 (100nM) and MST 312 (250nM). BIBR 1532 is a mixed-type non-competitive inhibitor, whereas MST 312 is a reversible inhibitor of telomerase activity (see Methods). Both inhibitors strikingly phenocopied two features of the *tert* mutant, in that we observed large early aggregate streams that broke and eventually resulted in mounds (Fig 4B; Table 3) and fruiting bodies that were small. The developmental delay, however, was not induced. Human TERT [48], which shares a 23% homology with *Dictyostelium* TERT, failed to rescue *tert* KO phenotype. Surprisingly, the morphologies of TERT overexpressing lines in the wild-type did not show any significant variation compared to those of the untreated wild-type (Fig 4A).

**Fig 4.**
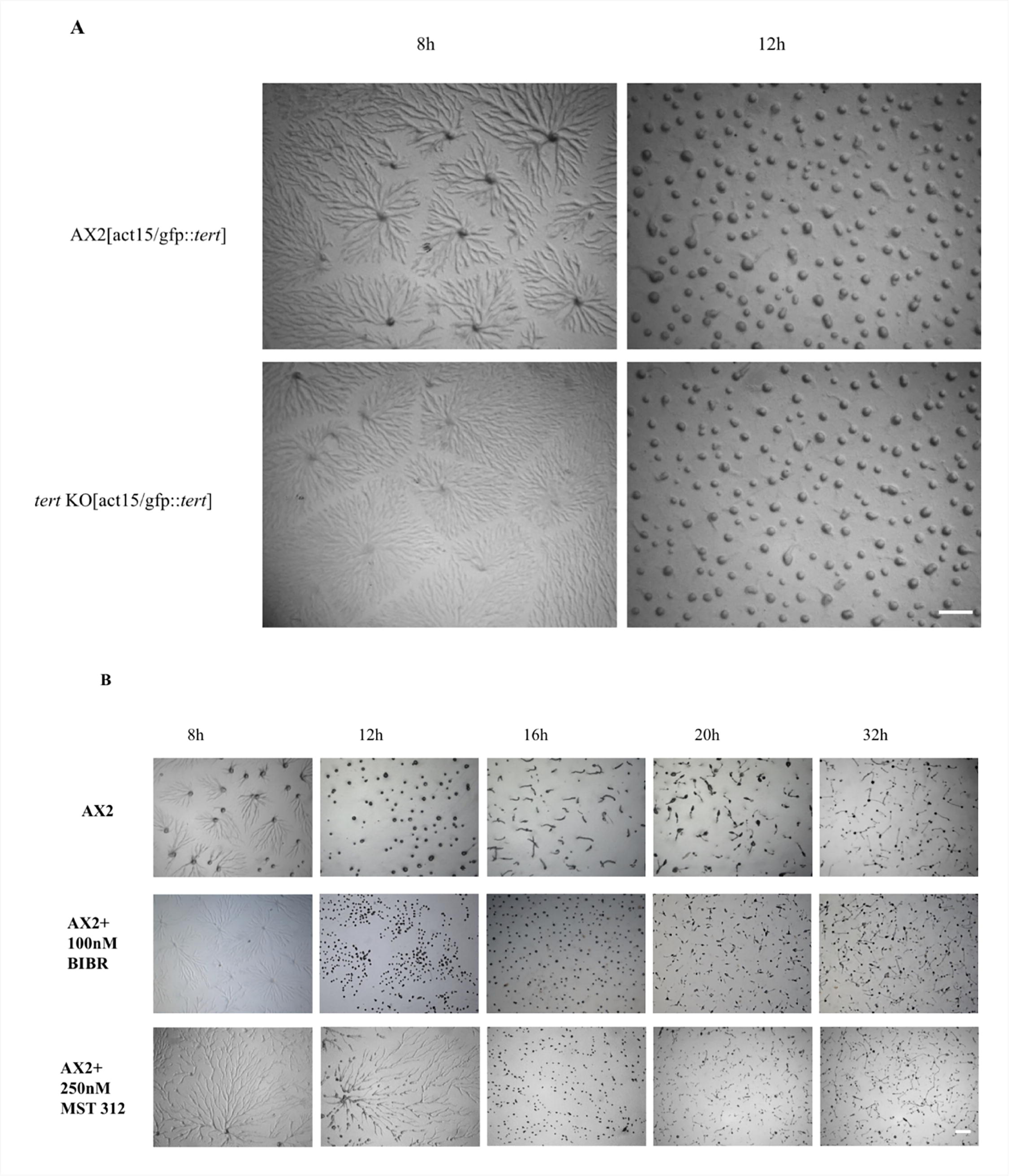
(A) Overexpression of TERT (act15/gfp::*tert)* rescued *tert* KO phenotype. Scale bar-0.5 mm. (B) AX2 cells treated with 100nM BIBR 1532 and 250 nM MST 312 phenocopied the *tert* KO streaming phenotype. The time points in hours are shown at the top. Scale bar-0.5 mm.

Overall, these results strongly support the idea that the relevant change in the *tert* KO involve *tert* itself. The fact that the TERT inhibitors induced only two of the three *tert* KO defects is interesting. Given the lack of any apparent interconnection between the pathway that regulates the switch to aggregation, and that regulating mound size, it seems likely that TERT acts on more than one molecular target. It could be that the inhibitors do not perturb that part of TERT that interacts with the target that regulates the switch to development.

### Roles of components of the mound size regulation pathway in the *tert* KO: *smlA, CF, countin* and glucose

#### smlA and countin

Compared to the wild-type, in the *tert* KO cells, *smlA* and *countin* expression levels were, respectively, low and high (Figs 5A and 5B; Table 1). Also, Western blots performed with anti-countin antibodies showed higher *countin* expression in *tert* KO cells, compared to wild-type (Fig 5C). When *tert* was overexpressed in the *tert* KO background, both *countin* and *smlA* expression levels were returned to those of the wild-type (Figs 5A and 5B). This overexpression also rescued all the defects of the *tert* KO phenotype (Fig 4A; Table 2). Given the previously proposed regulatory relationship between *smlA* and *countin* (Fig 1; [26,28,30]), the most parsimonious explanation for the majority of the results reported so far in this study, is that one role of *tert* in *D. discoideum* is to promote the expression of *smlA*, thus indirectly inhibiting *countin* expression, thus reducing glucose levels and mound/fruiting body size. This would make *tert* the most upstream regulator of these structures reported to date.

**Fig 5.**
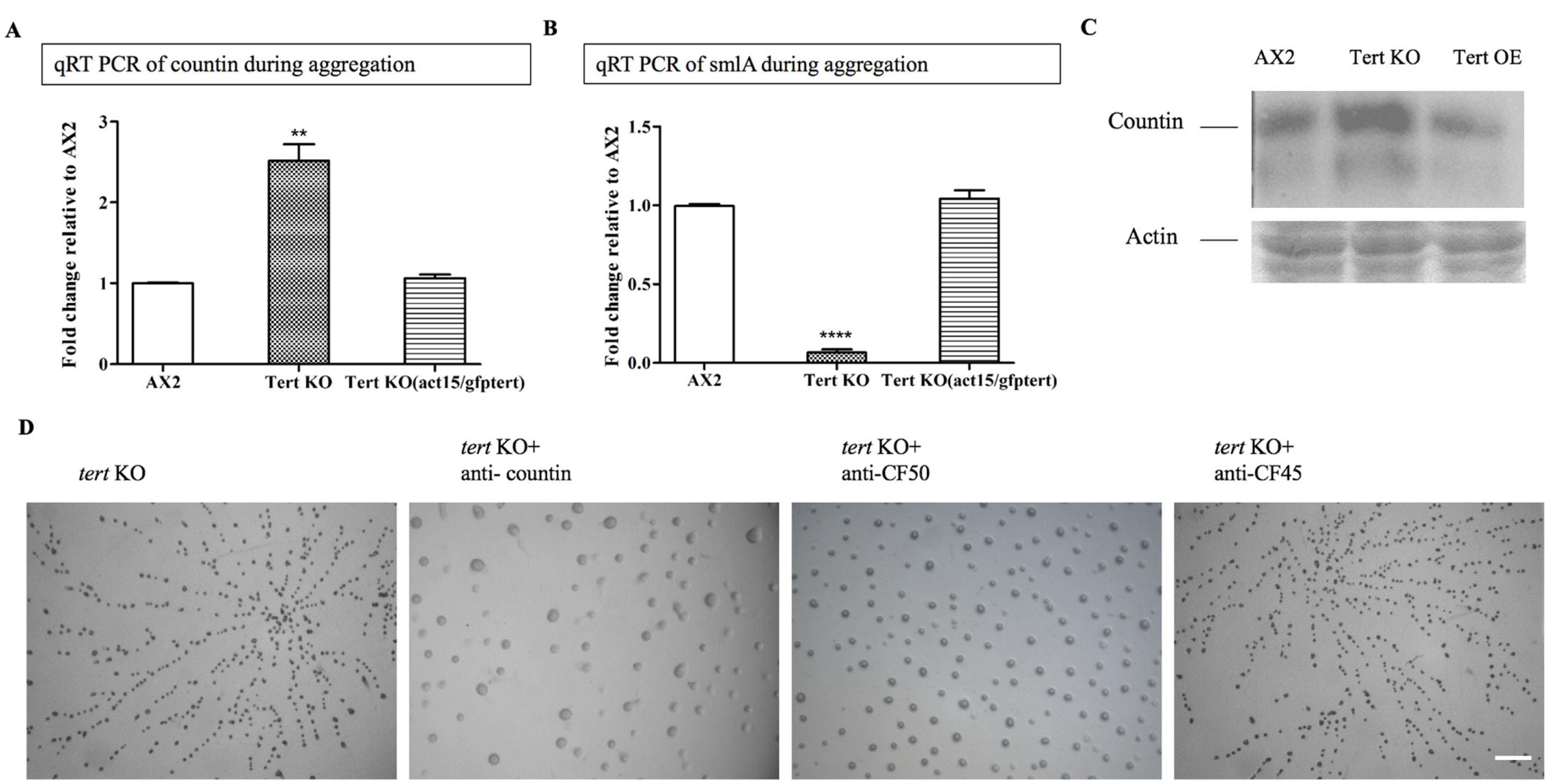
*Tert* regulates the levels of CF. qRT-PCR of (A) *countin* and (B) *smlA* during aggregation in AX2, *tert* KO and *tert* KO [act15/gfptert]. *rnlA* was used as mRNA amplification control. Level of significance is indicated as *p<0.05, **p<0.01, ***p<0.001, and ****p<0.0001. (C) Western blots with anti-countin antibodies. The gels were stained with Coomassie to show equal loading. (D) *Countin* KO cells were developed in the presence of *tert* KO conditioned media or BIBR1532. Scale bar- 0.5 mm. D) Cells were starved and developed with anti-countin, CF50, CF45, AprA and CfaD antibodies (1:300 dilution). Addition of anti-countin and anti-CF50 antibodies rescued *tert* KO group size defect. Scale bar- 0.5 mm.

The likelihood of some involvement of CF itself was supported by the effects of antibodies that target its components. When *tert* KO cells were treated with anti-countin or anti-CF50 antibodies (1:300 dilution), there was a reduction in aggregate fragmentation resulting in larger mounds compared to untreated *tert* KO controls (Fig 5D; Table 2); the development delay was not rescued. Adding anti-CF45 antibodies did not rescue any of the defects (Fig 5D; Table 2).

Indirect evidence that *tert* is acting upstream of CF was seen in the lack of effect of adding BIBR 1532 to *countin* KO cells, which typically exhibit no stream breaking and large mounds [28]. While, as noted above, BIBR 1532 leads to stream breaking and small mounds in wild-type cells, it did not lead to any change in the usual phenotype of *countin* KO cells (e.g. Fig 6A), which argues against *tert* acting downstream of *countin*.

**Fig 6.**
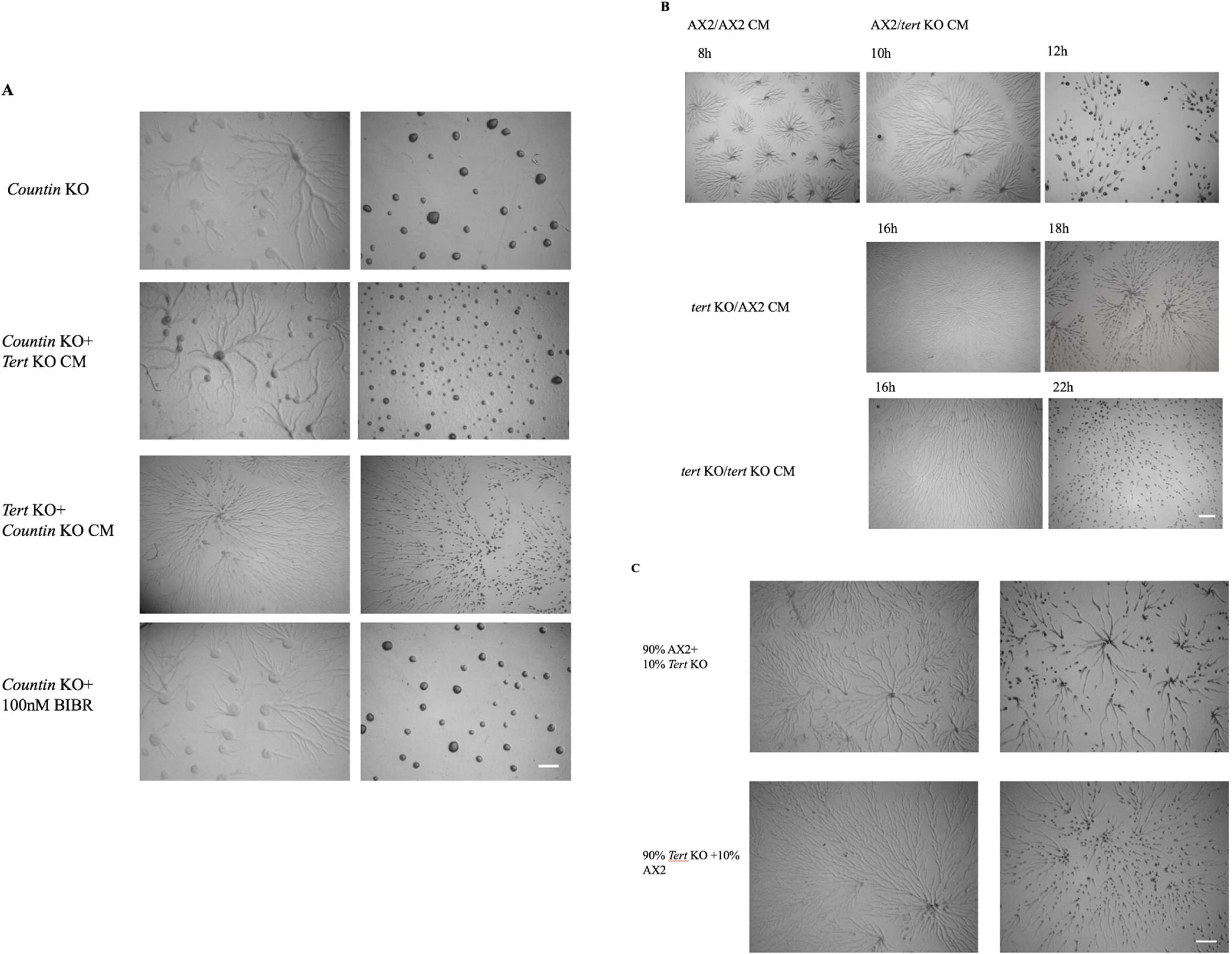
*Tert* regulates the levels of CF. (A) *Countin* KO cells were developed in the presence of *tert* KO conditioned media or BIBR1532. Scale bar- 0.5 mm. (B) Development in the presence of conditioned medium. *Tert* KO-CM induced stream breaking in AX2. (C) Reconstitution of AX2 in 1:9 ratio with *tert* KO did not rescue the stream breaking. Scale bar- 0.5 mm.

Beyond the observations already noted, a range of other observations support the idea that some of the *tert* KO’s features are due to the increased activity of a secreted mound-size regulating factor, such as countin. Conditioned medium (CM) from *tert* KO cells induced stream breaking in the wild-type (Fig 6B; Table 3) and led to reduced mound size. Also, adding *tert* KO CM to the *tert* KO itself aggravated the fragmentation phenotype (Fig 6B; Table 2). *Tert* KO CM was even capable of inducing stream fragmentation (Fig 6A), and reducing mound size, in *countin* mutants, suggesting that the CF levels of the *tert* KO CM were high. In each of these three cases, the *tert* KO CM not only affected streaming and mound size, but also induced, or aggravated, a development delay (Figs 6A and 6B; Tables 2 and 3). This suggests that the unknown TERT-induced factor that affects the developmental switch is also secreted.

Further, the presence of *tert* KO cells, even at very low concentrations (10%), was able to partially induce the *tert* KO phenotype when added to a population of wild-type cells and plated at an overall density of 5×10^5^ cells/cm^2^ (Fig 6C; Table 3). The apparent potency of the presumed high CF levels produced by the *tert* KO cells might partly explain one otherwise unexpected observation: Adding wild-type CM to *tert* KO cells did not rescue any of their defects (Fig 6B; Table 2). While the wild-type CM in this case would be expected to act as a diluent of CF (and thus potentially rescue the *tert* KO), this effect would only be brief. Development occurs over many hours, during which time the *tert* KO conditions could allow the build-up of CF back to mound-size-limiting levels. Similar reasoning might also explain why CM from *countin* KO cells (which exhibit undelayed aggregation and normal streaming) did not rescue any of the defects of *tert* KO cells (Fig 6A; Table 2).

We also observed if *tert* KO CM affected the wild-type cells of other Dictyostelids, such as *D. minutum, D. purpureum, D. fasciculatum* and *Polysphondylium pallidum* (each representing a distinct group in the Dictyostelid taxonomy). CM of *tert* KO did not affect the mound size of any of the species tested (Fig 7; Table 3) suggesting that some of the factors regulating mound size may be species specific.

**Fig 7.**
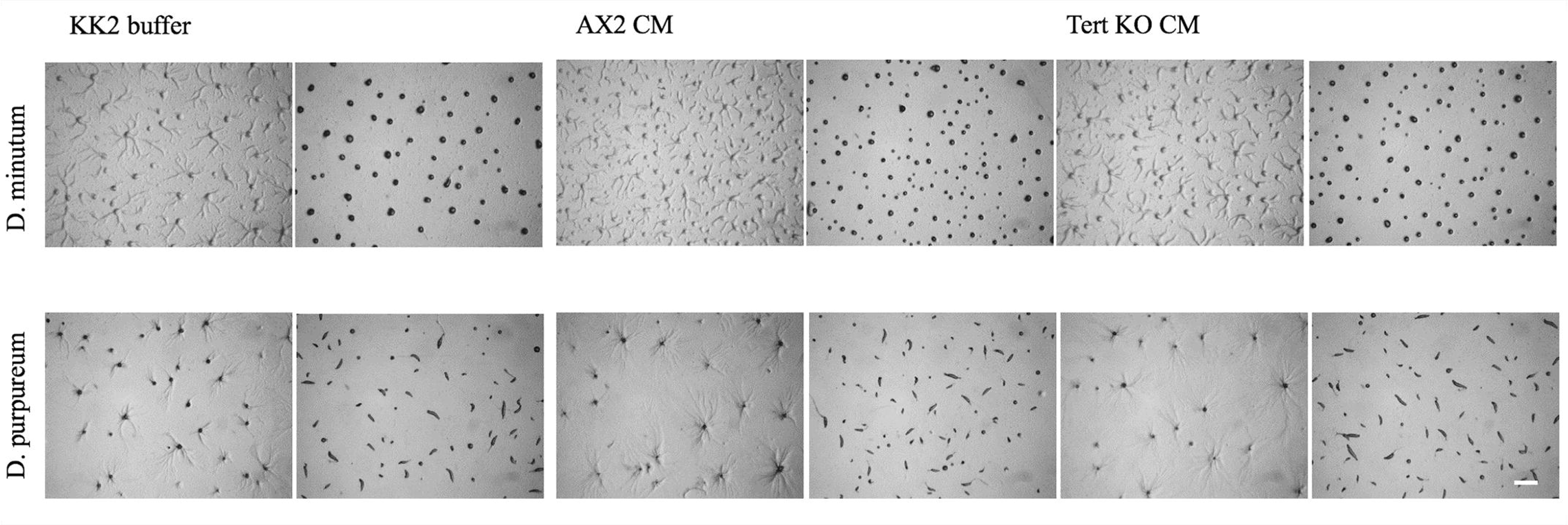
Development of other Dictyostelid species in the presence of *tert* KO conditioned medium. *tert* KO-CM did not alter the group size of other Dictyostelids. Scale bar- 0.5 mm.

#### Glucose rescued streaming and mound size defects, but not the delay

As per the model shown in Fig 1, one of the more downstream effects that should be seen if the *tert* KO has higher levels of CF, is the lowering of glucose levels. Glucose levels during aggregation were measured and in the *tert* KO were significantly lower (10.7±0.6 mg/ml) compared to wild-type (15.5±0.94 mg/ml) (Fig 8A, p=0.0015). Supplementing 1mM glucose rescued the aggregate streaming (and mound size), defects of the *tert* KO (Fig 8B), but not, as expected, the delay (Table 2).

**Fig 8.**
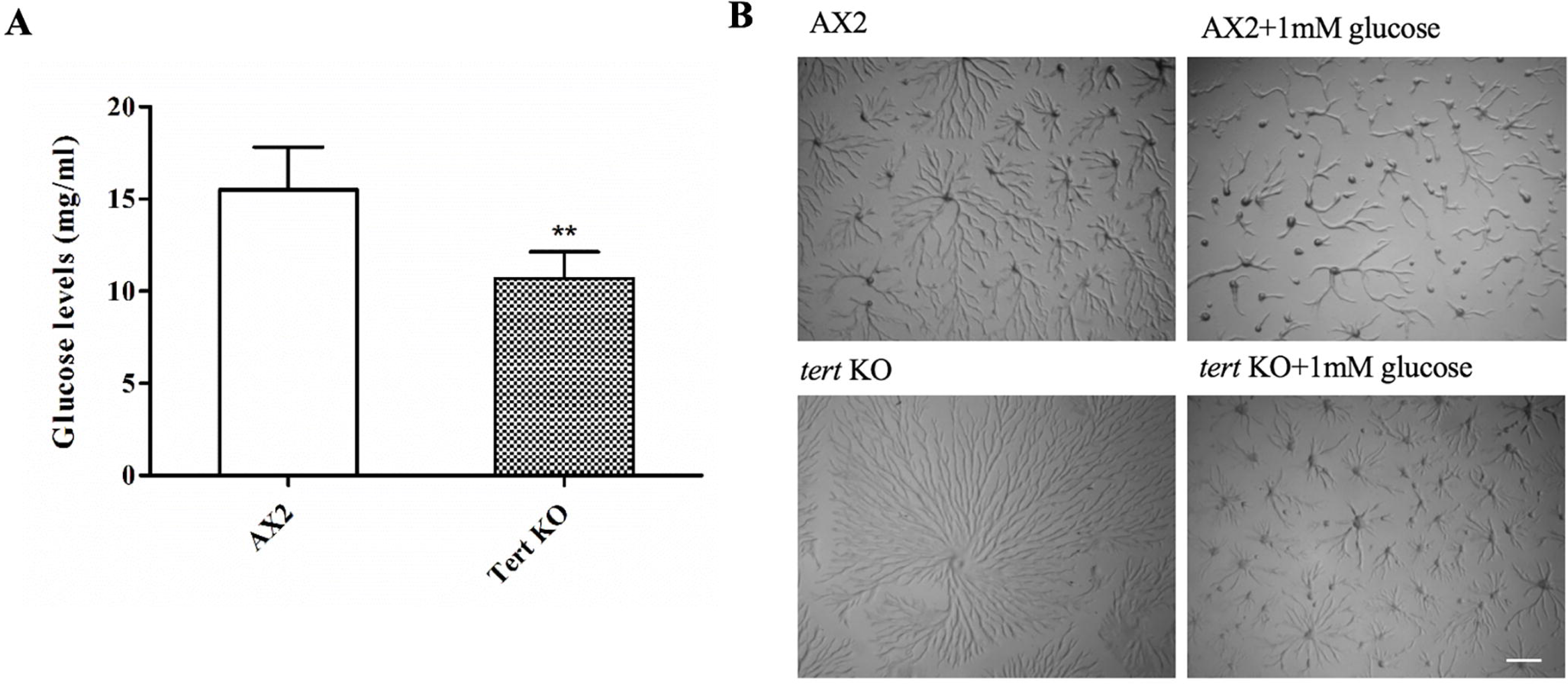
Effect of glucose on *tert* KO aggregate size. (A) Glucose levels during aggregation. (B) Wild-type AX2 and *tert* KO cells were developed in the presence of 1 mM glucose. Glucose rescues the streaming defect of *tert* KO. Scale bar- 0.5 mm. Level of significance is indicated as *p<0.05, **p<0.01, ***p<0.001, and ****p<0.0001.

#### Antibodies against AprA and CfaD did not rescue the tert KO phenotype

Previous work has shown that deletion of AprA and CfaD genes, involved in a different cell-density sensing pathway to that involving *smlA* and countin, leads to changes in mound-size [29], but, here, antibodies against AprA and CfaD did not rescue the KO phenotype (Fig 9), suggesting, again, that impaired mound size determination in the *tert* KO is largely due to defective CF signal transduction.

**Fig 9.**
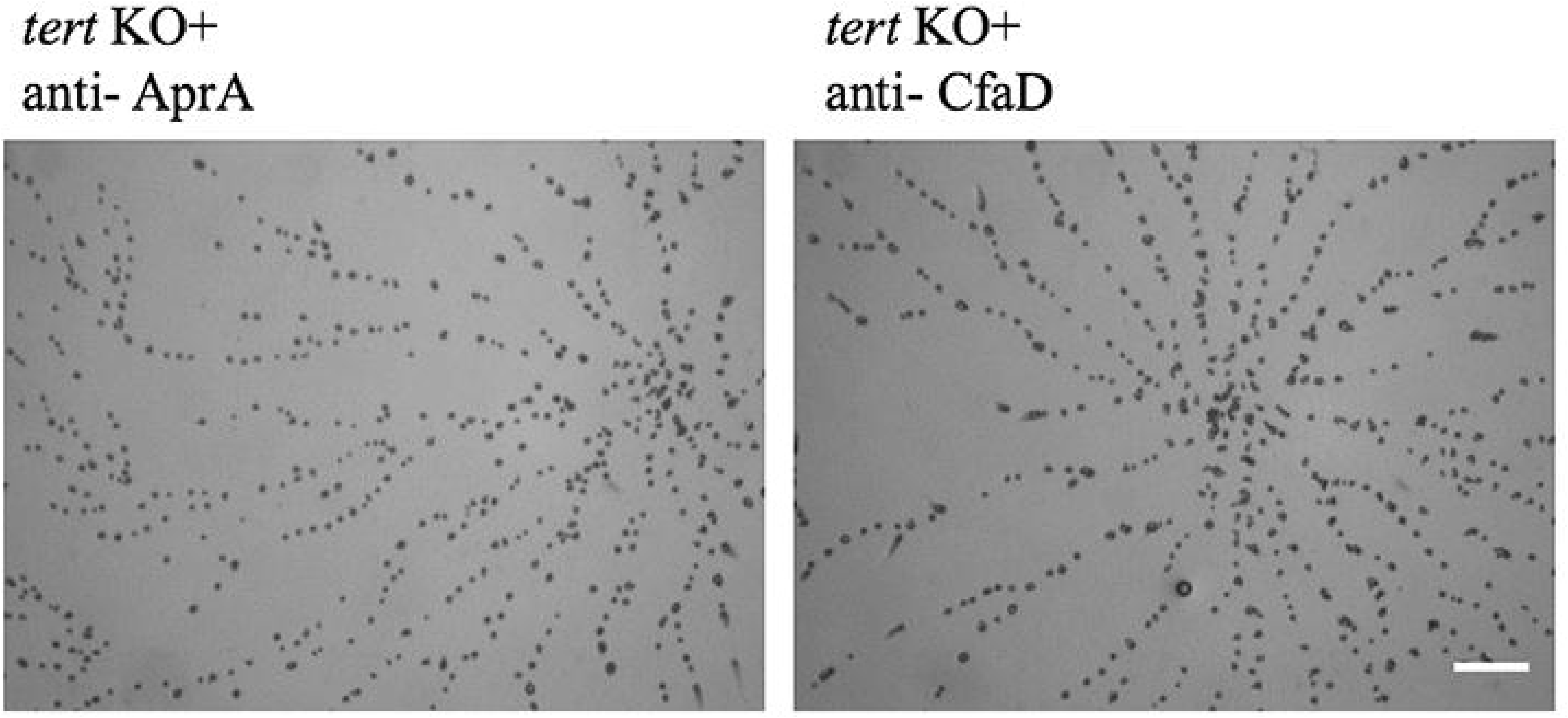
Cells were starved and developed with AprA and CfaD antibodies (1:300 dilution). Scale bar- 0.5 mm.

### Roles of cAMP and cAMP-related processes and factors in the *tert* KO

Given the perturbations seen in the *tert* KO, one would predict some abnormalities associated with cAMP dynamics [49–54]. The role of cAMP in streaming, in particular, has been much studied. Below we examine how various cAMP processes or factors, related to streaming and developmental delay, were affected in the *tert* KO.

#### Multiple cAMP wave generating centres observed in the tert KO

Starving cells normally aggregate by periodic synthesis and relay of cAMP, resulting in the outward propagation of cAMP waves from the aggregation centres [55]. We visualized cAMP waves by recording the time-lapse of development and then subtracting the image pairs [56]. Coordinated changes in cell shape and movement of cAMP waves can indirectly be visualized by dark field optics because of the differences in the optical density of cells moving/not moving in response to cAMP. Compared to the wild-type, which had a single wave generating centre, the *tert* KO had multiple wave propagating centres (Fig 10, Supplementary Videos 4 and 5). When the *tert* KO was rescued by overexpression of wild-type *tert*, so was the single wave propagating centre. The optical wave density analysis suggests that cAMP wave propagation is defective in *tert* KO, also contributing to stream breaking.

**Fig 10.**
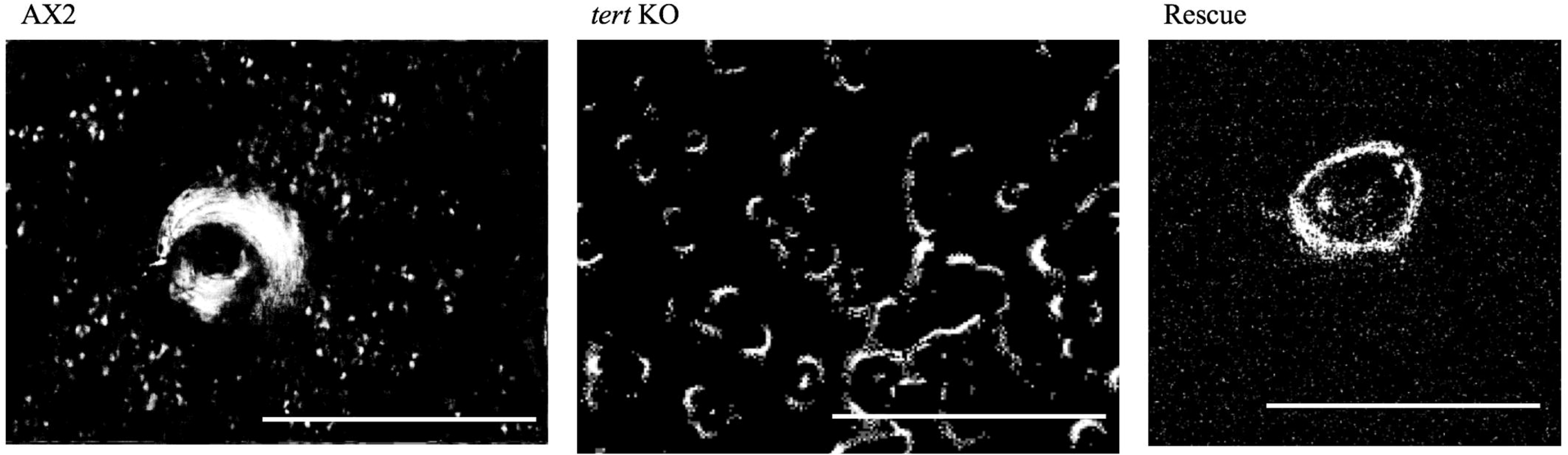
cAMP wave generating centres. Optical density wave images depicting wave generating centres in AX2, *tert* KO and rescue strain are shown. AX2 and rescue strain has a single wave generating centre, whereas *tert* KO has multiple wave generating centres in a single aggregate. Scale bar- 1 mm.

#### cAMP-related gene expression, cAMP levels, chemotaxis and relay were also impaired in the tert KO

Both the switch to aggregation, and normal streaming, require that a great variety of other cAMP-related processes occur properly. We quantified the relative expression of genes involved in cAMP synthesis and signaling in wild-type and *tert* KO cells by qRT-PCR during streaming as well as breaking (Figs 12A and 12B). With respect to the switch to aggregation, the expression levels of *acaA* (cAMP synthesis), *carA* (cAMP receptor), *5’NT* (5’ nucleotidase), *pdsA* (cAMP phosphodiesterases), *regA* and *pde4* were low initially but most started to ‘recover’ closer to the time that the *tert* KO manages to overcome its developmental delay (Figs 11A-F). Another, perhaps more meaningful, approach is to compare the levels in the mutant and wild-type when they are at equivalent developmental stages. This was done at two stages (aggregation, stream breaking) for four of the cAMP genes (*acaA, carA, pdsA, pde4*). During aggregation (i.e. at 8 h in the wild-type; 16 h in the *tert* KO), *acaA* and *carA* expression levels were significantly lower in the mutant, and the other two genes trended lower (Fig 12A). During stream breaking (10 h; 18 h, respectively), only *acaA* was significantly lower. Correspondingly, at 8 h of development, cAMP levels were also lower in the *tert* KO (0.98±0.08 nM in the KO; 1.59±0.15 nM in wild-type; Fig 12C, p=0.005). By 12 h, however, as the *tert* KO cells are closer to the time when their streaming will begin (i.e. 16 h) both cAMP-related gene expression, and cAMP levels increase, implying that the initially down-regulated expression of cAMP signaling might explain the long-delayed switch to aggregation in the *tert* KO. As to how cAMP-related genes or processes do recover in the absence of TERT, there are no indications in our results, but regulatory networks are well-known to exhibit a degree of robustness [57,58].

**Fig 11:**
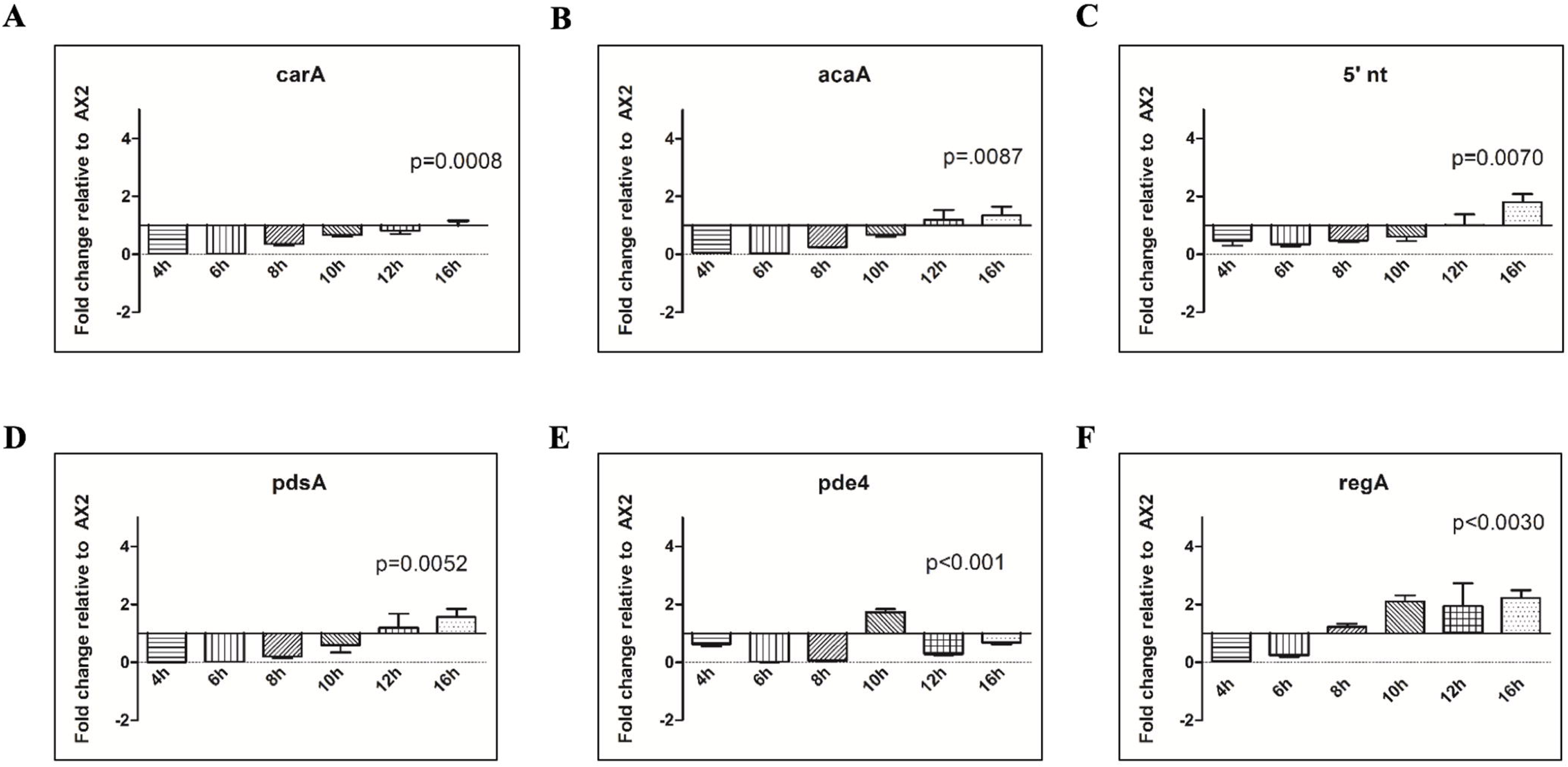
Delayed development in *tert* KO. (A-F) qRT-PCR of genes involved in the cAMP relay. Down-regulation of genes involved in the cAMP relay in *tert* KO. Fold change in mRNA expression is relative to AX2 at the indicated time points. *rnlA* is used as mRNA amplification control.

**Fig 12.**
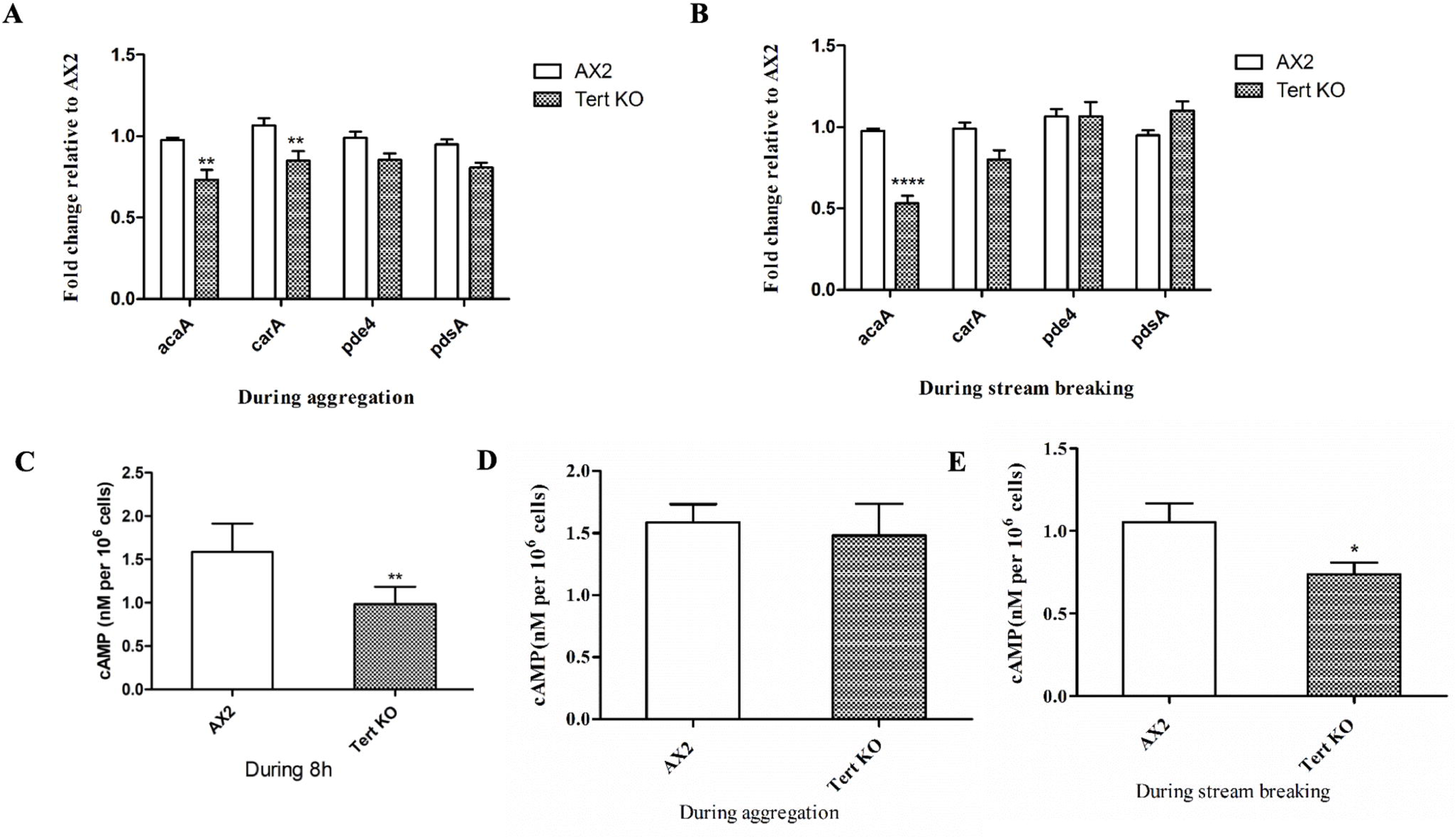
Defective cAMP relay of *tert* KO. cAMP relay and expression of *acaA, carA, pde4, pdsA* in *tert* KO during (A) aggregation and (B) stream breaking. Fold change in mRNA expression is relative to AX2 at the indicated time points. *rnlA* was used as mRNA amplification control. cAMP levels in *tert* KO during (C) 8 h of development in AX2 and *tert* KO, (D) aggregation, (E) stream breaking. Level of significance is indicated as *p<0.05, **p<0.01, ***p<0.001, and ****p<0.0001.

As noted, cAMP-related gene expression levels of the *tert* KO lag behind that of the wild-type, and even though they increase as the mutant enters a similar developmental phase, the cAMP levels never catch up completely. When cAMP levels were quantified during aggregation and stream breaking using an ELISA-based competitive immunoassay, the cAMP levels in the wild-type and *tert* KO were 1.59±0.15 nM and 1.48±0.25 nM, respectively, during aggregation (Fig 12D, p=0.73); and 1.05±0.11 nM and 0.74±0.70 nM during stream breaking (Fig 12E, p=0.04). Thus, these lower absolute levels of cAMP in the *tert* KO may also contribute to abnormal stream breaking, with the amoebae unable to relay signals to their neighbours.

To test whether cAMP-based chemotaxis was normal, we performed an under-agarose chemotaxis assay, towards 10 µM cAMP. The trajectories of cells were tracked and their chemotaxis parameters were quantified. Although the speed of cells towards cAMP was higher in *tert* KO (16.01±1.39 µm/min) compared to the wild-type (12.74±0.43 µm/min), the directionality was significantly reduced in *tert* KO cells (0.37±0.03 compared 0.59±0.04). The chemotactic index of *tert* KO cells also was lower (0.63±0.05) compared to wild-type cells (0.82±0.06) (Figs 13A-C).

**Fig 13.**
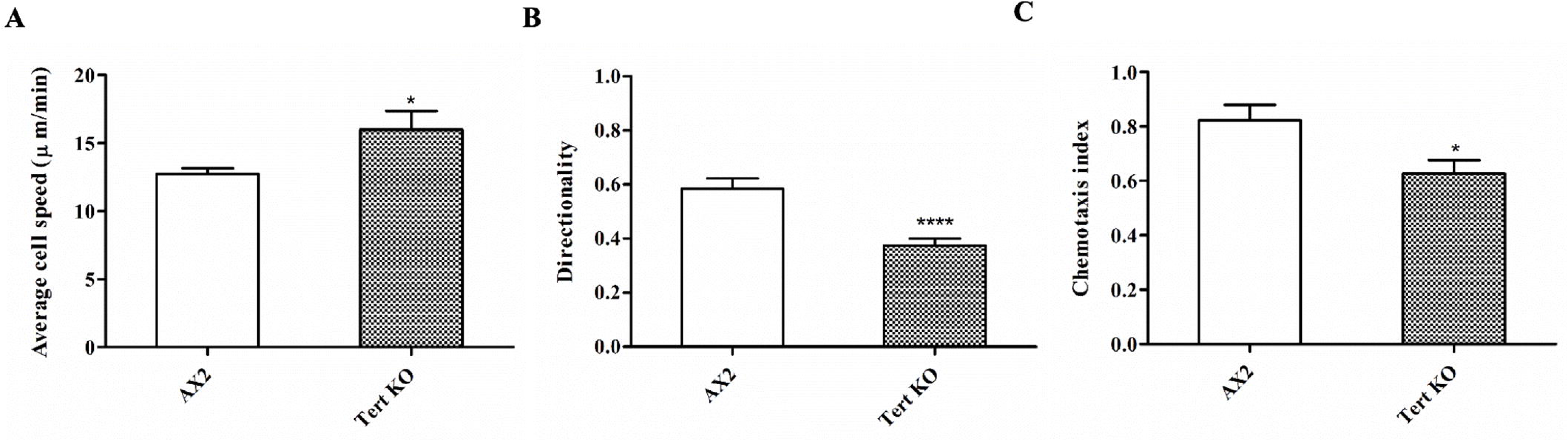
Defective cAMP chemotaxis of *tert* KO. Under-agarose cAMP chemotaxis assay in response to 10µM cAMP. (A) Average chemotaxis speed in response to cAMP. (B) directionality of chemotaxing cells and (C) chemotaxis index are shown. The graph represents the mean and SEM of 3 independent experiments.

#### The chemotaxis defect of the tert KO was not rescued by cAMP pulsing or 8-Br-cAMP

To gain further insights into the streaming defect of the *tert* KO cells, we examined if cAMP pulsing could rescue the chemotaxis defect [59,60]. cAMP (50nM) pulsing was carried out every 6 minutes for 4 hours and thereafter, the cells were seeded in the starvation buffer at a density of 5×10^5^ cells/cm^2^ and different developmental stages were monitored (Fig 14A). The streaming defect of *tert* KO was not rescued by cAMP pulsing, suggesting that other components of cAMP signaling are necessary to rescue the defect.

**Fig 14.**
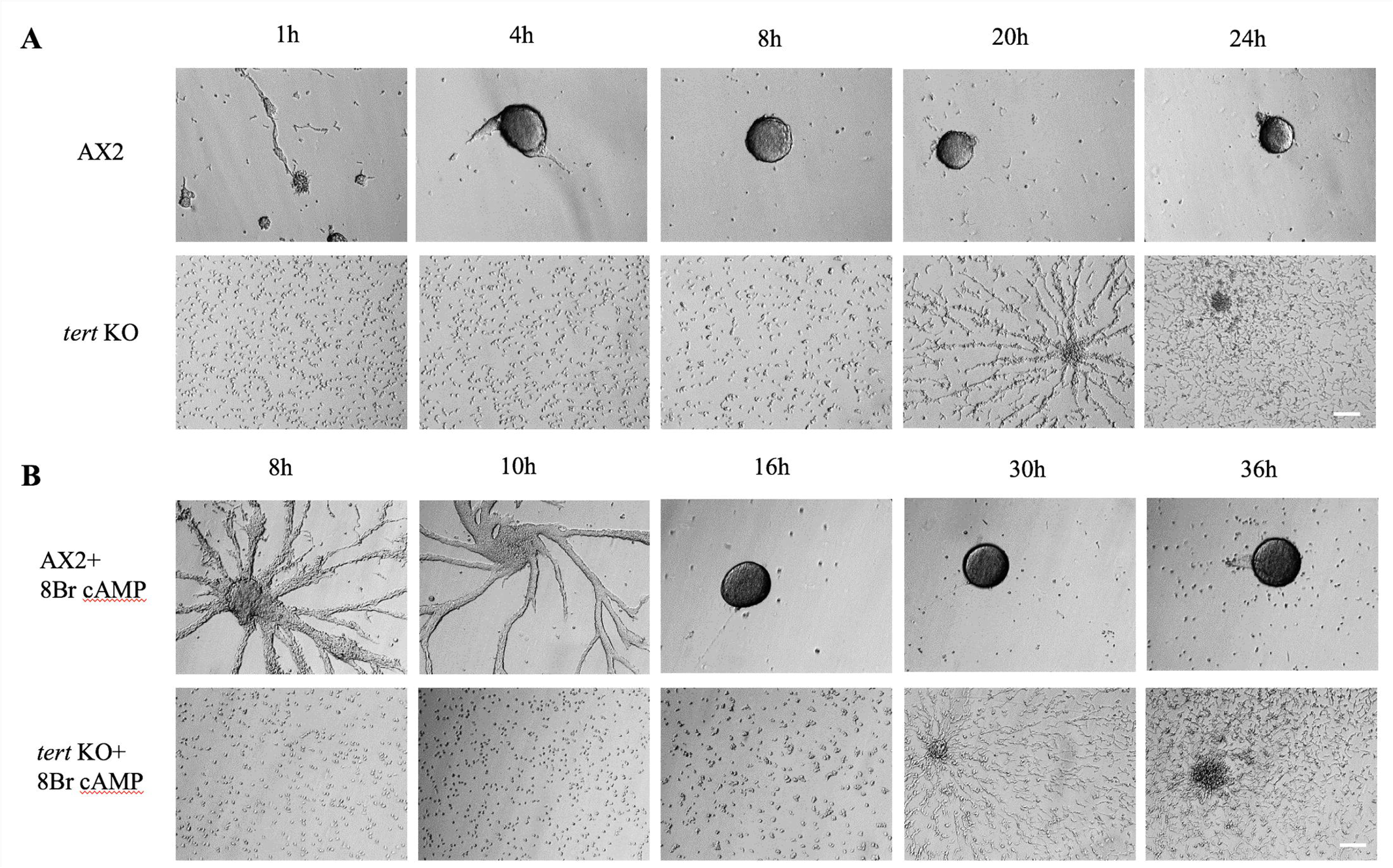
cAMP sensing in *tert* KO. (A) Wild-type and *tert* KO cells were starved for 1 hour and pulsed every 6 min with 50 nM cAMP for 4 h. Cells were then resuspended in BSS and seeded at a density of 1×10^5^ cells/ml, and observed under a microscope. (B) Wild-type and *tert* KO cells were washed in BSS, seeded at a density of 1×10^5^ cells/ ml, and incubated in BSS or BSS + 5 mM 8-Br-cAMP for 5 h. Cells were washed and then observed under a microscope. Scale bar- 100 µm.

If cAMP receptor activity is compromised, that could also lead to defective signaling and to test this, we used a membrane-permeable cAMP analog 8-Br-cAMP, which has poor affinity for extracellular cAMP receptors and enters the cells directly [61]. Cells were incubated with 5mM 8-Br-cAMP and after 5 h, the cells were transferred to Bonners Salt Solution and different developmental stages were monitored (Fig 14B). Adding 8-Br-cAMP did not rescue the *tert* KO phenotype, suggesting that cAMP receptor function is not impaired in the mutant.

#### High adenosine levels in the tert KO induced large aggregation streams

As mentioned previously, adenosine and caffeine are known to alter the cAMP relay [62,63], thereby affecting aggregate size. This occurs in a number of Dictyostelids [33]. We observed enhanced expression of *5’NT* in the *tert* KO (Fig 15A, p=0.0042) suggesting increased adenosine levels (5’NT converts AMP to adenosine). Hence, adenosine levels were quantified and these were significantly higher (235.37±26.44 nM/10^6^ cells) in *tert* KO cells compared to wild-type (35.39±12.78 nM/10^6^ cells) (Fig 15B, p=0.0051). The adenosine antagonist, caffeine (1 mM), rescued the streaming defect (Fig 15C), and restored the mound size, suggesting that excess adenosine in the *tert* KO causes larger streams. It did not, however, rescue the developmental delay. Since glucose also rescues the streaming defect in *tert* KO cells, adenosine levels were quantified subsequent to treating with 1mM glucose. Glucose treatment reduced adenosine levels (13.07±7.51 nM/10^6^ cells) in *tert* KO cells to a level that is more comparable to wild-type cells (35.39±12.78 nM/10^6^ cells), but as already noted, it did not rescue the developmental delay. Importantly, *5’NT* expression and adenosine levels reduced significantly subsequent to stream breaking (S4 Fig). This could perhaps be due to negative feedback on *tert* itself. When wild-type cells were treated with 2mM adenosine, *tert* expression levels were reduced, although not significantly so (S5 Fig).

**Fig 15.**
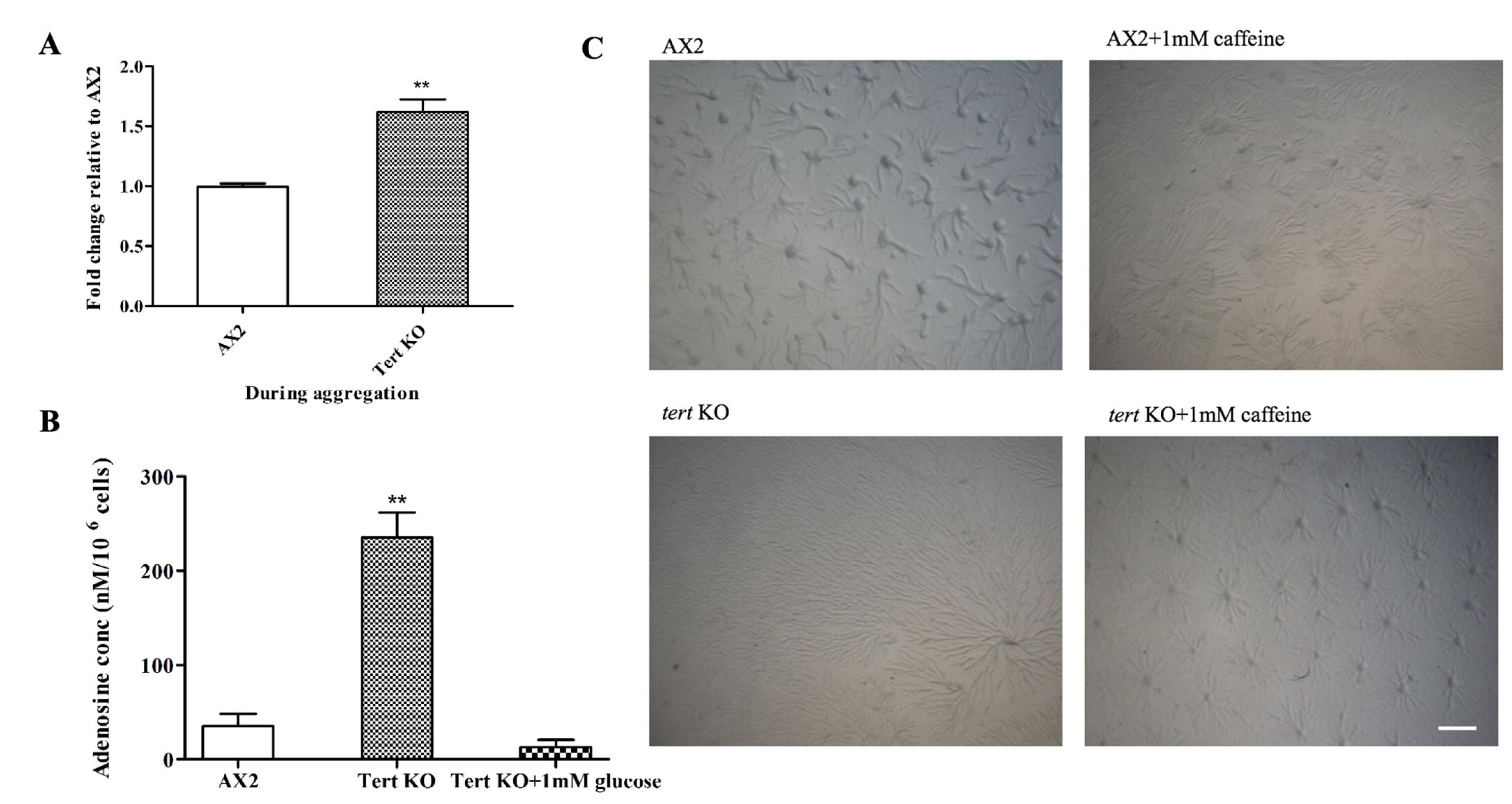
Effect of adenosine on aggregate size. (A) qRT-PCR of 5’NT. Fold change in mRNA expression is relative to AX2 at indicated time points. *rnlA* is used as mRNA amplification control. (B) Quantification of adenosine levels. Level of significance is indicated as *p<0.05, **p<0.01, ***p<0.001, and ****p<0.0001. (C) Cells were developed in the presence of 1mM caffeine; *tert* KO streaming defect was rescued. Scale bar- 0.5 mm.

#### Streaming defects of the tert KO were not due to increased iron levels

*Dictyostelium* cells, when grown in the presence of 200µM iron, formed large streams that fragmented into multiple mounds, strikingly resembling the *tert* KO phenotype [64]. As the phenotypes had similarities, we examined if TERT mediates its effect by altering intracellular iron levels. We quantified iron by ICP-MS and the levels were not significantly different between the wild-type (16.38±1.21 ng/10^7^ cells) and *tert* KO cells (15.25±0.81 ng/10^7^ cells) (S6 Fig, p=0.4573), suggesting that *tert* KO phenotype is not due to altered iron levels.

### The role of adhesion-related factors in the *tert* KO, as they affect streaming and mound size

Cell-substratum adhesion is also important for migration and proper streaming. By spinning cells at different speeds (0, 25, 50 and 75 rpm), it is possible to vary substratum dependent sheer force. Thus, by counting the fraction of floating cells at different speeds, it is possible to check substratum dependent adhesion. Although both wild-type and *tert* KO cells exhibited a sheer force-dependent decrease in cell-substratum adhesion, *tert* KO cells exhibited a significantly weaker cell-substratum adhesion (Fig 16, p<0.0001), thus indicating a contribution to stream breaking.

**Fig 16.**
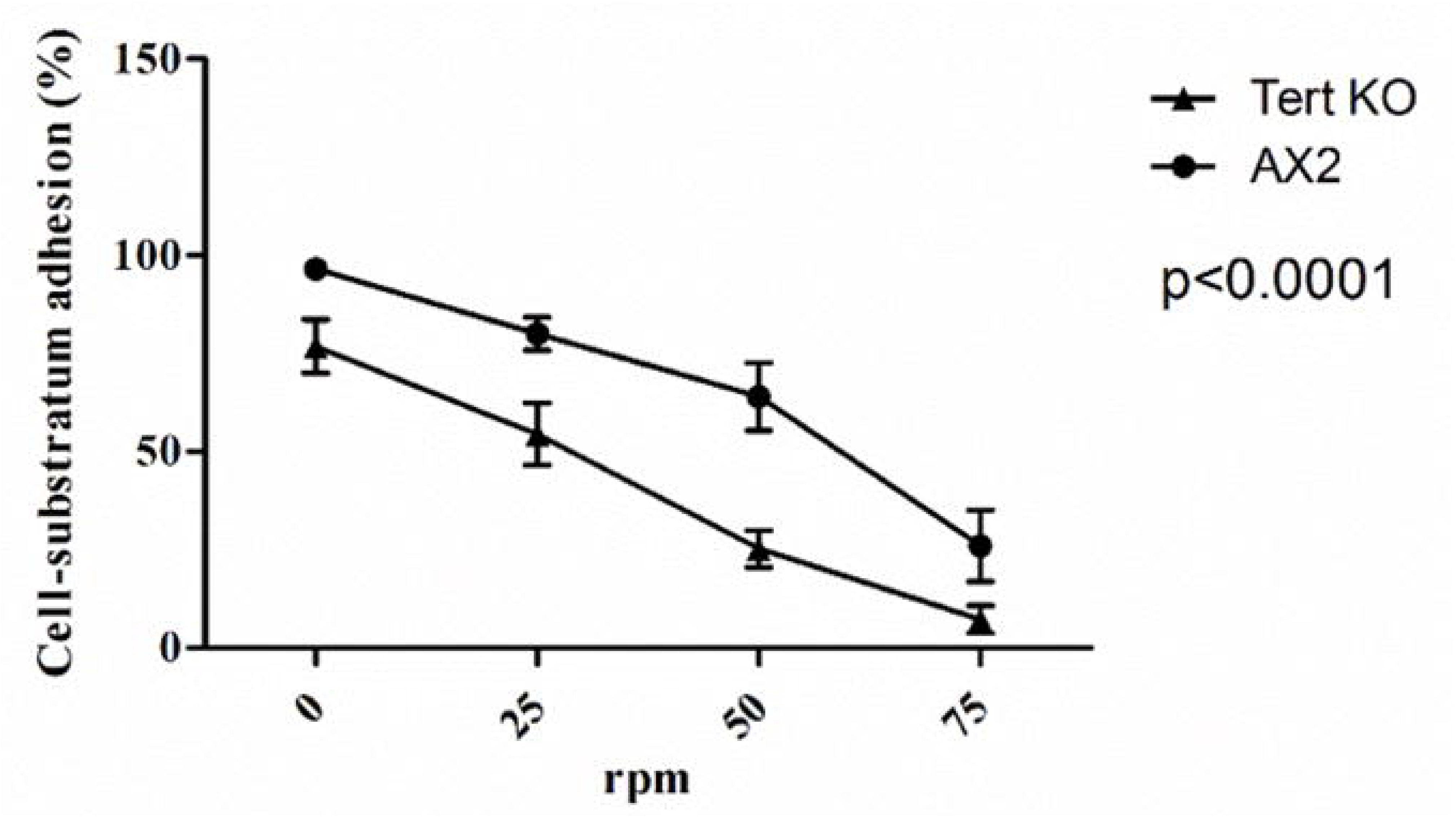
Disruption of *tert* affects cell substratum adhesion. Cells were plated at a density of 1×10^5^ cells/ml, grown overnight, in an orbital shaker. Floating and attached cells were counted and percentage adhesion was plotted versus rotation speed. Both AX2 and *tert* KO exhibited a sheer force-dependent decrease in substratum adhesion and *tert* KO exhibited significantly reduced adhesion compared to AX2 cells.

Cell-cell adhesion is also an important determinant of streaming and mound size in *Dictyostelium* [39]. To examine if adhesion is impaired in the mutant, we checked the expression of two major cell adhesion proteins, *cadA*, expressed post-starvation (2 h) and *csaA* expressed during early aggregation (6 h). *cadA*-mediated cell-cell adhesion is Ca^2+^-dependent and thus EDTA-sensitive, while *csaA* is Ca^2+^ independent and EDTA-resistant [65]. Both *csaA* and *cadA* expression were significantly down-regulated (Figs 17A and 17B).

**Fig 17.**
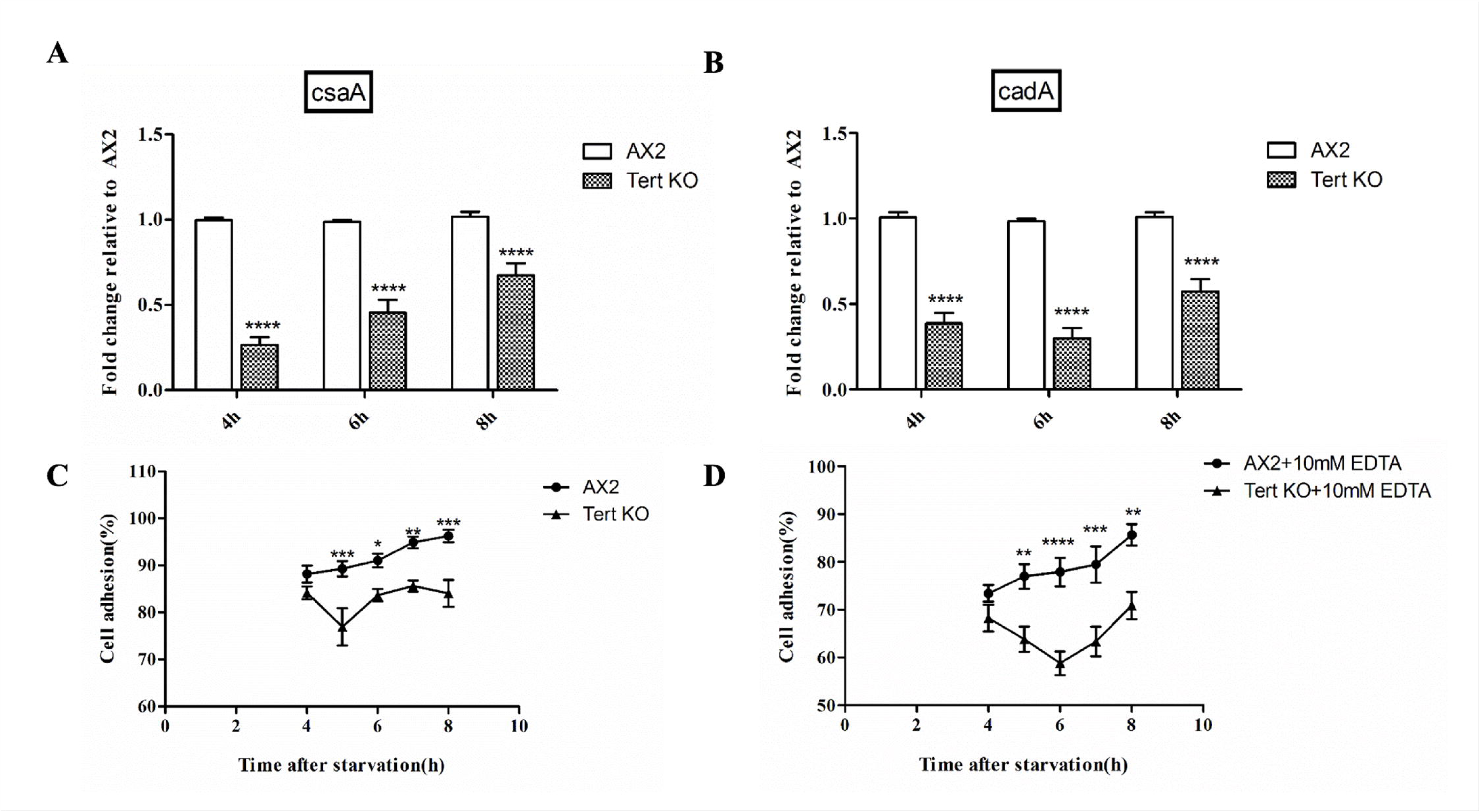
Disruption of *tert* affects cell adhesion. qRT-PCR of (A) *csaA* and (B) *cadA. rnlA* was used as mRNA amplification control. Wild-type and *tert* KO cells were starved in Sorensen phosphate buffer at 150 rpm and 22 ºC. Samples were collected at the start of the assay and at one-hour time points after 4 h of starvation. Percentage of cell adhesion plotted over time. (C) EDTA resistant cell-cell adhesion, (D) EDTA sensitive cell-cell adhesion. Level of significance is indicated as *p<0.05, **p<0.01, ***p<0.001, and ****p<0.0001.

Further, cell adhesion was monitored indirectly by counting the fraction of single cells not joining the aggregate. Aggregation results in the gradual disappearance of single cells and thus it is possible to measure aggregation by determining the ratio of single cells remaining. To examine Ca^2+^-dependent cell-cell adhesion, the assay was performed in the presence of 10 mM EDTA. Both EDTA-sensitive and resistant cell-cell adhesion were significantly defective in *tert* KO cells (Figs 17C, p=0.0033 and 17D, p=0.0015).

These results imply that defective cell-substratum, and cell-cell adhesion might play roles in the abnormal streaming and mound-size regulation of the *tert* KO.

### The developmental delay of the *tert* KO was associated with reduced polyphosphate levels

One interesting observation was that the only treatment that fully rescued the *tert* KO cells was the overexpression of wild-type *tert*. Also, the only other treatment that rescued the developmental delay itself was mixing wild-type cells with the *tert* KO cells at a 1:1 ratio (Fig 18; Table 2). Even though caffeine and glucose rescued streaming and mound size, and apparently this was at least partly mediated via their impact on cAMP-regulated processes, neither of the compounds rescued the delay, even though abnormalities of cAMP-regulated processes are commonly reported causes of delay in other *Dictyostelium* studies [52–54].

**Fig 18.**
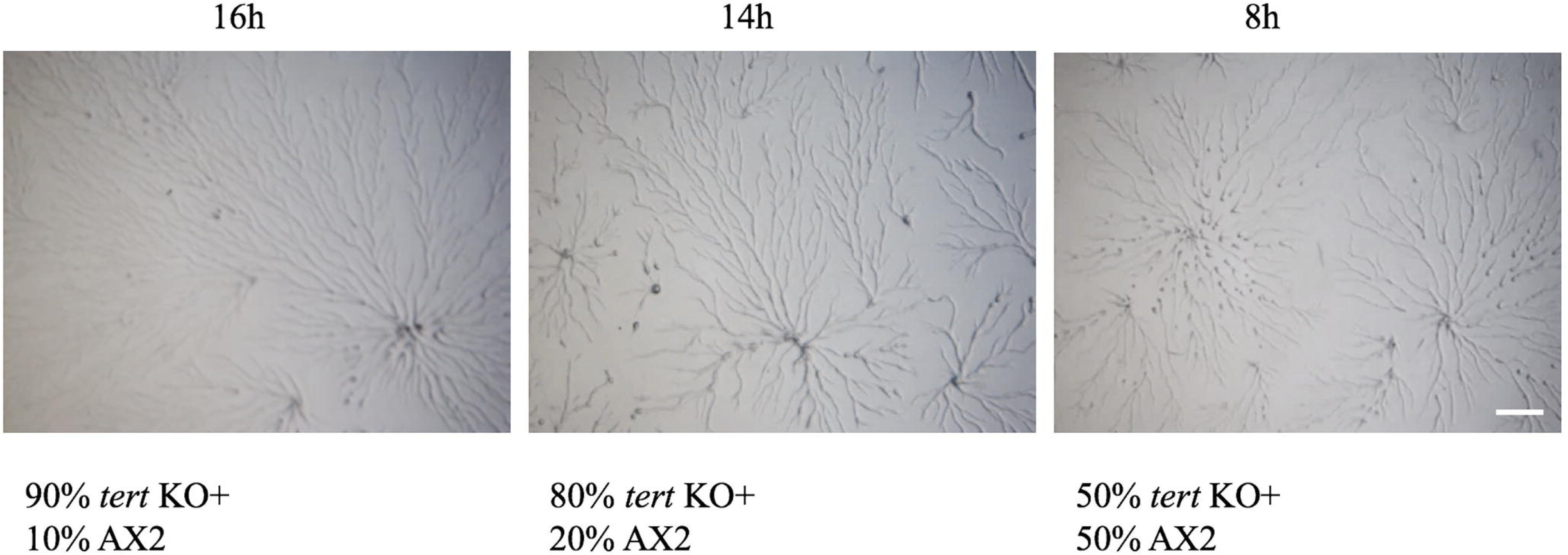
Rescue of delay by added wild-type cells. Wild-type AX2 and *tert* KO were reconstituted at 1:9, 2:8 and 1:1 ratio. Developmental delay of *tert* KO was rescued by AX2 at 1:1 ratio. Scale bar- 0.5 mm.

Thus, we examined polyphosphate levels in the *tert* KO because of their known importance to developmental timing in *Dictyostelium* [66]. We stained the CM with DAPI for 5 mins and checked the polyphosphate specific fluorescence using a spectrofluorometer. The CM of *tert* KO cells has reduced polyphosphate levels (49.55±2.02 µM) compared to wild-type (60.62±1.95 µM), implying that low polyphosphate levels might also contribute to the delay in initiating development in this system (Fig 19, p=0.0009).

**Fig 19.**
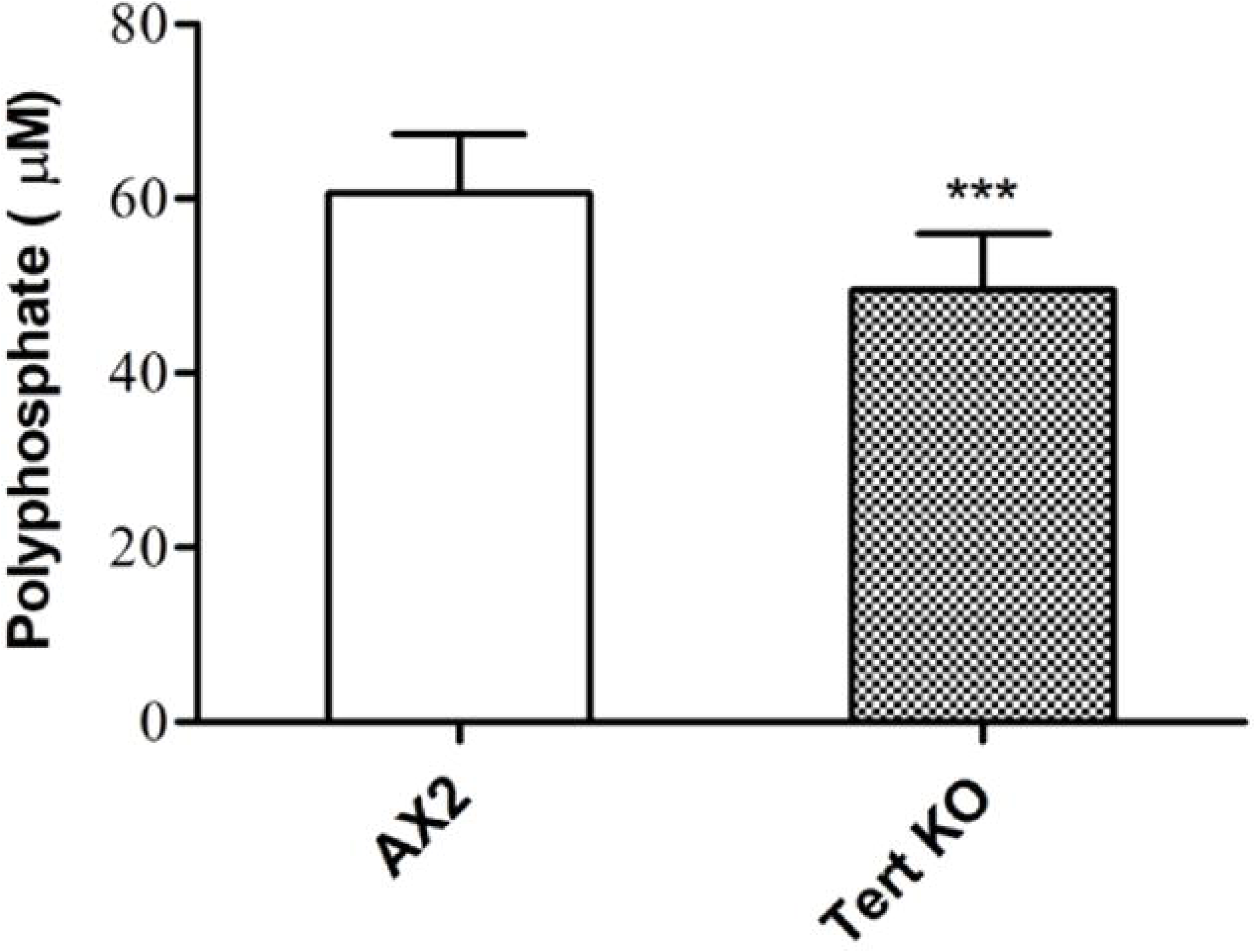
Polyphosphate levels were low in the *tert* KO. Polyphosphate levels in conditioned media of AX2 and *tert* KO. Level of significance is indicated as *p<0.05, **p<0.01, ***p<0.001, and ****p<0.0001.

## Conclusions

Our results reveal that TERT plays an important role in many aspects of *Dictyostelium* development. The *tert* KO exhibited a wide range of developmental defects. Despite suitable environmental conditions for multicellular development to begin, the start of the streaming phase is delayed by 8 h. Having once begun, development proceeds and ends abnormally, with large streams, uneven fragmentation, and, eventually, small mounds and fruiting bodies. The wide-ranging developmental defects are associated with changes to the levels, or expression, of genes and compounds that are known to be highly upstream regulators of the various stages of development, such as streaming and mound/fruiting body formation. Based on the perturbations in the *tert* KO, and our other experiments, Fig 20 depicts the possible extent of processes, and potential mediating factors, that might depend upon normal *tert* expression/TERT activity in the wild-type. Note that the arrows that connect *tert*/TERT to any element in the diagram are not meant to suggest that TERT directly regulates that element, only that TERT is important, perhaps in some indirect way, for the normal levels, or activity, of that element.

**Fig 20.**
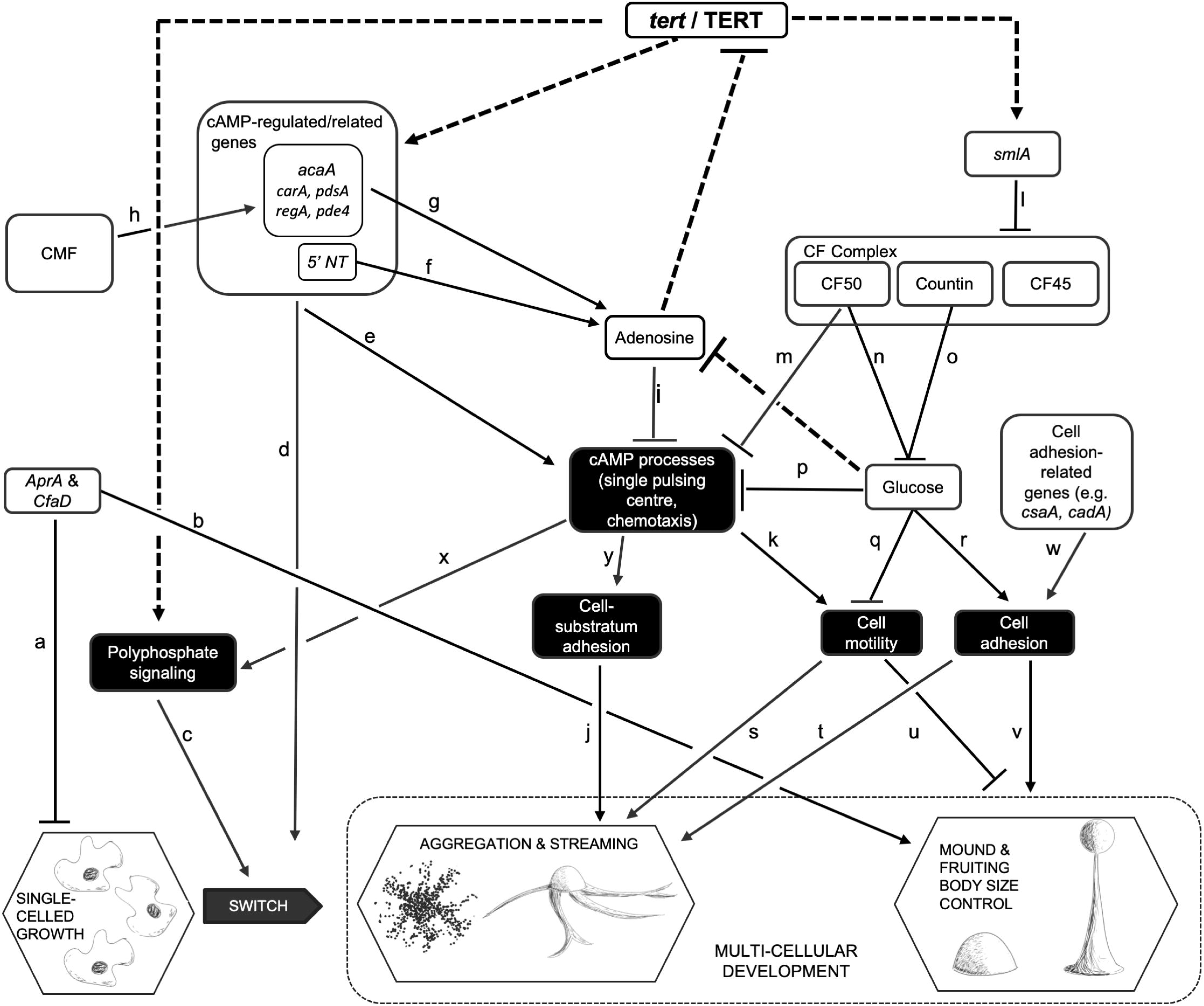
Some of the possible targets of *tert*/TERT in development of *D. discoideum,* as indicated by this study. This work, the first functional study of a telomerase in *Dictyostelium*, revealed that TERT influenced many previously reported developmental processes and pathways. The dashed lines represent effects previously unreported, involving multiples phases of the life-cycle. Adenosine, however, was found to provide negative feedback on *tert* expression. The depictions of life-cycle changes were drawn by the authors, based on images in [102], and adapted by kind permission of the Journal of Cell Science.

One of the most striking findings was that TERT appears to regulate, or is at least necessary for, the normal activity of what was previously known as the most upstream regulator of mound size, *smlA* [26,28,30]. Expression levels of *smlA* were reduced in the *tert* KO, and we also observed a wide variety of the expected downstream effects of lowered *smlA* levels. All of these, and a wide variety of treatments that rescued the size-defect of the mutant phenotype, support the idea that the reduction of mound size in the *tert* KO was indeed mediated via the abnormal functioning of the previously-identified elements of the mound-size regulation pathway.

In addition to the rescue approach, treatments that attempted to phenocopy the *tert* KO phenotype in the wild-type, also suggest TERT is a highly upstream regulator of mound size. In particular, given that size regulation in *D. discoideum* depends upon secreted factors of the CF complex, one would have predicted the effects we observed when *tert* KO CM was added to wild-type cells. Another strong indication that *tert* acts upstream, at least of CF, was that the inhibition of *tert* activity in *countin* mutants failed to phenocopy the *tert* KO phenotype.

A similarly rich range of results (involving the *tert* KO phenotype, and its rescue, and phenocopying) support the idea that TERT also plays a high-level role in the regulation of streaming. During the streaming phase, two genes associated with cAMP related-processes in *D. discoideum* (*acaA, carA*) were significantly downregulated (compared to the wild-type), and the levels of several other genes trended lower. This was also accompanied by lower cAMP levels. This could explain the defective chemotaxis and cell motility of the *tert* KO.

Of course, the regulation of streaming is not entirely isolated from that of size. Glucose, one of the central elements of the CF pathway, influences several cAMP-related processes [67]. Thus, it was not surprising that adding 1mM glucose to the *tert* KO cells rescued both the size and streaming defects. This study, however, provided a new insight into how the rescue of streaming occurs, because added glucose also reduced adenosine levels. Thus, in the *tert* KO, the low glucose levels might lead to higher adenosine levels, allowing it to inhibit cAMP related processes (via pathway i, Fig 20). In normal development, given that the known sequence of the telomere repeats of *D. discoideum* (A-G_(1-8)_; [43]), and the fact that telomerase activity would therefore recruit cellular stores of adenosine, it is possible that normal TERT activity keeps adenosine levels low. As yet, however, whether TERT actually acts as a functional telomerase in *D. discoideum* is not known.

The *tert* gene we characterized includes the conserved domains and structure of a telomerase reverse transcriptase. Also, supplementing structurally unrelated but specific inhibitors of TERT to wild type cells phenocopies the mutant phenotype. The widely used method to test telomerase activity is the TRAP assay. However, this method failed to detect telomerase activity in *D. discoideum* and there may be both technical and innate limitations. For example, possible reasons are that: (i) the presence of rDNA palindrome elements in the chromosomal ends, suggesting a novel telomere structure and the possible role of TERT in maintaining both rDNA and chromosomal termini; and (ii) polyasparagine repeats, present in the TERT protein of *Dictyostelium,* splitting the functional domain into two halves. It is not clear whether the loss of TERT function is due to the absence of normal eukaryote telomeres in *D. discoideum* [68].

The discussion so far, while it establishes that TERT is needed for several developmental processes to take place, does not help to distinguish whether or not it acts more than once, or if it has more than one target. Could TERT for example act more like the much studied homeodomain proteins, master regulators of animal development, but which only act during very early embryological life [69,70]? Likewise, in *D. discoideum,* CMF appears to act only once [32]. Two lines of argument suggest that TERT is different.

First, the biphasic nature of *tert*’s expression pattern suggest that it acts during at least two stages of development. In the wild-type, *tert* expression builds up to its first peak at 8 h, thus being a strong candidate for enabling streaming to begin, and to proceed correctly, around this time. It then dips markedly to a low point at 10 h, whereby it might help to enable stream break-up by its relative absence. Then, it begins its climb to its second peak at 12 h, when mound size is being finalised.

Second, while it is well known that cAMP-related processes play important roles in allowing streaming to begin and to proceed properly, and while we have shown that TERT influences multiple cAMP related processes, the pathway by which TERT influences the initiation of streaming seems distinct from that used for maintaining it. Both glucose and caffeine, for example, rescued the streaming and size defects of the *tert* KO, but the delay was unaffected. Complementarily, when wild-type cells were mixed at 50% with *tert* KO cells, they rescued the delay defect only. In fact, the only treatment that fully rescued the *tert* KO was the overexpression of wild-type *tert.*

This study indicates for, the first time, that TERT acts in several non-canonical ways in *D. discoideum,* influencing when aggregation begins, the processes involved in streaming, and the eventual size of the fruiting body. TERT’s influences appear to occur upstream of many other regulators, and, in particular, TERT could be the most upstream regulator of mound and fruiting body size identified so far. Curiously, as yet we have no evidence that TERT acts as a canonical telomerase, nor is it known whether any other enzyme protects the unusually sequenced telomeres of this species. In the most heavily studied stages of the organism’s life-cycle, that is, those that occur in response to starvation, replication has ceased, so further study of this particular point should focus on the amoeboid stage. More generally, this study has revealed several previously unreported non-canonical processes influenced by a telomerase. TERT’s roles in influencing cell motility and adhesion, and the levels of chalone-like secreted factors, bear consideration by those engaged in cancer research.

## Methods

### *Dictyostelium* culture and development

Wild-type *D. discoideum* (AX2) cells were grown with *Klebsiella aerogenes* on SM/5 plates, or axenically, in modified maltose-HL5 medium (28.4 g bacteriological peptone, 15 g yeast extract, 18 g maltose monohydrate, 0.641 g Na_2_HPO_4_ and 0.49 g KH_2_PO_4_ per litre, pH 6.4) containing 100 units penicillin and 100 mg/ml streptomycin-sulphate. Cells were also grown in Petri dishes as monolayers. Other Dictyostelid species (*D. minutum, D. purpureum, D. facsiculatum* and *Polysphondylym pallidum*) were grown with *Klebsiella aerogenes* on SM/5 plates and cells were harvested when there was visible clearing of bacterial lawns.

To trigger development, cells were washed with KK_2_ buffer (2.25 g KH_2_PO_4_ and 0.67 g K_2_HPO_4_ per liter, pH 6.4) and plated on 1% non-nutrient KK_2_ agar plates at a density of 5×10^5^ cells/cm^2^ in a dark, moist chamber [71]. To study streaming, cells were seeded in submerged condition (KK_2_ buffer) at a density of 5×10^5^ cells/cm^2^.

BIBR 1532 is a specific non-competitive inhibitor of TERT with IC50 value of 93 nM for human telomerase [72]. To find the optimal dose response of BIBR 1532 in Dictyostelium, starved cells were plated in phosphate buffered agar with different concentrations of BIBR 1532 (10 nM, 25 nM, 50 nM, 100nM and 200 nM) and 100nM was found to be the minimal effective dose in inducing complete stream breaking. MST 312, which is structurally unrelated to BIBR 1532, is a reversible inhibitor of TERT with IC50 value of 0.67 µM for human telomerase [73]. The minimal effective dose in *Dictyostelium* was found to be 250 nM.

### Construction of *tert* expression vector

Using genomic DNA as template, a 3.8kb *tert* sequence was PCR amplified using ExTaq polymerase (Takara) and ligated in pDXA-GFP2 vector by exploiting the HindIII and KpnI restriction sites. This vector was electroporated to *tert* KO and AX2 cells and G418 resistant (10 µg/ml) clones were selected and overexpression was confirmed by semi-quantitative PCR. Primer sequences used for generating the vectors are mentioned in S1 Table.

### Conditioned medium assay

Conditioned medium was prepared as described previously with slight modifications [74]. Briefly, log phase cells of AX2 and *tert* KO were resuspended at a density of 1×10^7^ cells/ml and kept under shaking conditions for 20 h. Cells were pelleted and the supernatant was further clarified by centrifugation. The clarified supernatant (CM) was used immediately. To check the effect of CM on aggregate size, cells were developed in the presence of CM on non-nutrient agar plates and development was monitored. KK_2_ buffer was used as control. To deplete extracellular CF with anti-countin antibodies, cells were starved in KK_2_ buffer. After 1 h, the cells were developed with anti-countin antisera (1:300 dilution) in KK_2_ [75].

### Western blot

To examine countin protein expression levels during aggregation, Western blot was performed with anti-countin antibody. Cells were resuspended in SDS laemmli buffer, and boiled for 3 min. Subsequently, the samples were run in a 12% SDS-polyacrylamide gel and Western blots were developed using an ECL Western blotting kit (Biorad). Rabbit anti-countin antibodies were used at 1: 3000 dilutions.

### Cell-cell adhesion assay

Log phase cells were starved at a density of 1×10^7^ cells/ml in KK_2_ buffer in shaking conditions at 22 °C for 4 h. At the beginning of starvation, 4×10^7^ cells were removed and resuspended in 2 mL Sorensen phosphate buffer, vortexed vigorously and 0.4 mL of cell suspension was pipetted immediately in vials containing 0.4 ml ice-cold Sorensen phosphate buffer or 0.4 mL of 20 mM EDTA solution. The cell suspension was then transferred to a shaker and incubated for 30 min and 0.2 mL of 10% glutaraldehyde was added to each sample at the end of incubation and stored for 10 min. Then, 7 ml Sorensen phosphate buffer was added to each vial. Cell adhesion was indirectly measured by counting the number of single cells left behind using a hemocytometer [76].

### Cell-substratum adhesion

To measure cell-substratum adhesion, 5×10^5^ cells were seeded in 60mm Petri dishes and incubated at 22 °C for 12 hours. The Petri dishes with the cell suspension was placed on an orbital shaker at different speeds (0, 25, 50, 75 rpm). After 1 h, adherent and non-adherent cells were harvested, counted using a hemocytometer and the fraction of adherent cells was plotted against the rotation speed [77].

### Visualization of cAMP waves

To visualize cAMP wave propagation, 5×10^5^ cells/cm^2^ were plated on 1% non-nutrient agar plates and developed in dark moist conditions at 22 °C. On a real-time basis, the aggregates were filmed at an interval of 30 s/frame, using a Nikon CCD camera and documented with NIS-Elements D software (Nikon, Japan). For visualizing cAMP optical density waves, image pairs were subtracted [56] using Image J (NIH, Bethesda, MD).

### Under agarose cAMP chemotaxis assay

The under agarose cAMP chemotaxis assay was performed as described previously [78]. Briefly, 100 µl of cell suspension starved at a density of 1×10^7^ cells/ml in KK_2_ buffer was added to outer troughs and 10 µM cAMP was added in the middle trough of a 1% agarose plate. Cells migrating towards cAMP was recorded every 30 s for 15 min with an inverted Nikon Eclipse TE2000 microscope using NIS-Elements D software (Nikon, Japan). For calculating the average velocity, directionality and chemotactic index, each time 36 cells were analyzed. The cells were tracked using ImageJ. Velocity was calculated by dividing the total displacement of cells with time. Directionality was calculated as the ratio of absolute distance traveled to the total path length, where a maximum value of 1 represents a straight path without deviations. Chemotactic index was calculated as the ratio of the average velocity of a cell moving against a cAMP gradient to the average cell speed. It is a global measure of direction of cell motion.

### Quantitative real-time PCR (qRT-PCR)

Total RNA was isolated from AX2 and *tert* KO cells at the indicated time points (0–24 h) using TRIzol reagent (Life Technologies, USA) [79]. RNA samples were quantified with a spectrophotometer (Eppendorf) and were also analyzed on 1% TAE agarose gels. cDNA was synthesized from total RNA using cDNA synthesis kit (Verso, Thermo-scientific). 1 µg of total RNA was used as a template to synthesize cDNA using random primers provided by the manufacturer. 1 µl of cDNA was used for qRT-PCR, using SYBR Green Master Mix (Thermo-scientific). qRT-PCR was carried out to analyze the expression levels of *tert, acaA, carA, pdsA, regA, pde4*, 5’NT, *countin* and *smlA* using the QuantStudio Flex 7 (Thermo-Fischer). *rnlA* was used as mRNA amplification control. All the qRT-PCR data were analyzed as described [80]. The primer sequences are mentioned in S2 Table.

### cAMP quantification

cAMP levels were quantitated using cAMP-XP TM assay kit as per the manufacturer’s protocol (Cell signalling, USA). AX2 and *tert* KO cells developed on 1% KK_2_ agar, were lysed with 100 µl of 1X lysis buffer and incubated on ice for 10 min. 50 µl of the lysate and 50 µl HRP-linked cAMP solution were added to the assay plates, incubated at room temperature (RT) on a horizontal orbital shaker. The wells were emptied after 3 hours, washed thrice with 200 µl of 1X wash buffer. 100 µl of tetramethylbenzidine (TMB) substrate was added and incubated at RT for 10 min. The reaction was terminated by adding 100 µl of stop solution and the absorbance was measured at an optical density of 450 nm. The cAMP standard curve was used to calculate absolute cAMP levels.

### Glucose quantification

Glucose levels were quantified as per the manufacturer’s protocol (GAHK20; Sigma-Aldrich). Mid-log phase cells were harvested and resuspended at a density of 8×10^6^ cells/ml in KK_2_ buffer and kept in shaking conditions at 22 °C. Cells were collected again and lysed by freeze-thaw method. 35 μl of the supernatant was mixed with 200 µl of glucose assay reagent and incubated for 15 minutes. The absorbance was measured at an optical density of 540 nm. The glucose standard curve was used to calculate absolute glucose levels.

### Adenosine quantification

Adenosine quantification was performed as per the manufacturer’s protocol (MET5090; Cellbio Labs). Cells grown in HL5 media was washed and seeded at a density of 5×10^5^ cells/cm^2^ on KK_2_ agar plates. The aggregates were harvested using the lysis buffer (62.5mM Tris-HCl, pH 6.8, 2% SDS, 10% glycerol). 50 µl sample was mixed with control mix (without adenosine deaminase) or reaction mix (with adenosine deaminase) in separate wells and incubated for 15 minutes. The fluorescence was measured using a spectrofluorometer (Ex-550nm, Em-595nm). The adenosine fluorescence in the sample was calculated by subtracting fluorescence of control mixed sample from reaction mixed sample. The adenosine standard curve was used to calculate absolute adenosine levels.

### Polyphosphate measurements

The conditioned media was incubated with 25 µg/ml DAPI for 5 minutes and polyphosphate specific fluorescence was measured using a spectrofluorometer (Ex-415nm, Em-550nm) as previously described [81]. Conditioned medium samples were prepared in FM minimal media to reduce the amount of background fluorescence. Polyphosphate concentration, in terms of phosphate monomers were determined using polyphosphate standards.

### Microscopy

A Nikon SMZ-1000 stereo zoom microscope with epifluorescence optics, Nikon 80i Eclipse upright microscope or a Nikon Eclipse TE2000 inverted microscope equipped with a digital sight DS-5MC camera (Nikon) were used for microscopy. Images were processed with NIS-Elements D (Nikon) or Image J.

### Statistical tools

Microsoft Excel (2016) was used for data analyses. Unpaired student’s t-test and two-way ANOVA (GraphPad Prism, version 6) were used to determine the statistical significance.

## Acknowledgments

We thank the Dictyostelium Stock Center, USA for supplying *Dictyostelium* strains and plasmids. We thank Dr Richard Gomer (Texas A&M University) for providing countin, CF50, CF45, AprA and CfaD antibodies. Polyphosphate standards were a kind gift from Dr Toshikazu Shiba (RegeneTiss Inc.). hTERT cDNA was a kind gift from Dr. Jayakrishnan Nandakumar (University of Michigan). The telomerase activity assay protocol was suggested by Dr Elizabeth Blackburn. NN acknowledges Rakesh Mani, Shalini Umachandran, Prajna A Rai and J Meenakshi for discussions.

## Author Contributions

NN, RB designed the experiments. NN performed the experiments. NN, GH and RB analyzed the data. NN, GH and RB wrote the manuscript and all authors read and approved the final manuscript.

## Supporting information

**S1 Fig. Schematic representation of the different functional domains of TERT identified with SMART analysis.** TERT protein contains the following domains: reverse transcriptase and RNA binding domain.

**S2 Fig. Telomerase activity assay.** TRAP assay was performed for AX2 and *tert* KO. Human cell lines HEK and HeLa were used as positive controls.

**S3 Fig. Changing cell density and its effect on development.** Development assay at different cell density (2×10^4^ cells/cm^2^ to 2×10^6^ cells/cm^2^). AX2 cells aggregate even at a cell density below 2×10^4^ cells/cm^2^, but *tert* KO fails to aggregate at such a density. *Tert* KO phenotype was not rescued even at higher cell density (2×10^6^ cells/cm^2^).

**S4 Fig. Effect of adenosine on aggregate size in *D. discoideum*.** A) qRT-PCR of 5’NT during stream breaking. Fold change in mRNA expression is relative to AX2 at the indicated time points. *rnlA* is used as mRNA amplification control. B) Quantification of adenosine levels during stream breaking. Level of significance is indicated as *p<0.05, **p<0.01, ***p<0.001, and ****p<0.0001.

**S5 Fig. *Tert* levels in adenosine treated cells.**

**S6 Fig. Quantification of iron.** Iron levels were quantified by ICP-MS. Level of significance is indicated as *p<0.05, **p<0.01, ***p<0.001, and ****p<0.0001.

**S1 Table.** Primers used for TERT overexpression vector construction.

**S2 Table.** Primers used for real-time PCR.

**Video 1.** Timelapse video of AX2 development.

**Video 2.** Timelapse video of *tert* KO development.

**Video 3.** Timelapse video of *tert* KO (act15/gfp::*tert*) development.

**Video 4.** Timelapse video of cAMP wave propagation in AX2.

**Video 5.** Timelapse video of cAMP wave propagation in *tert* KO.

