## Supplementary information for "A telomerase with novel non-canonical roles: TERT controls cellular aggregation and tissue size in *Dictyostelium*"

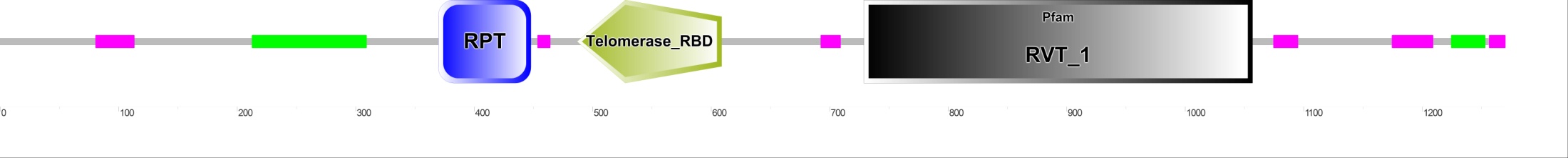

**S1 Fig**

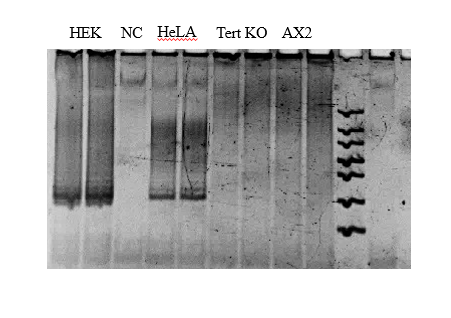

**S2 Fig**

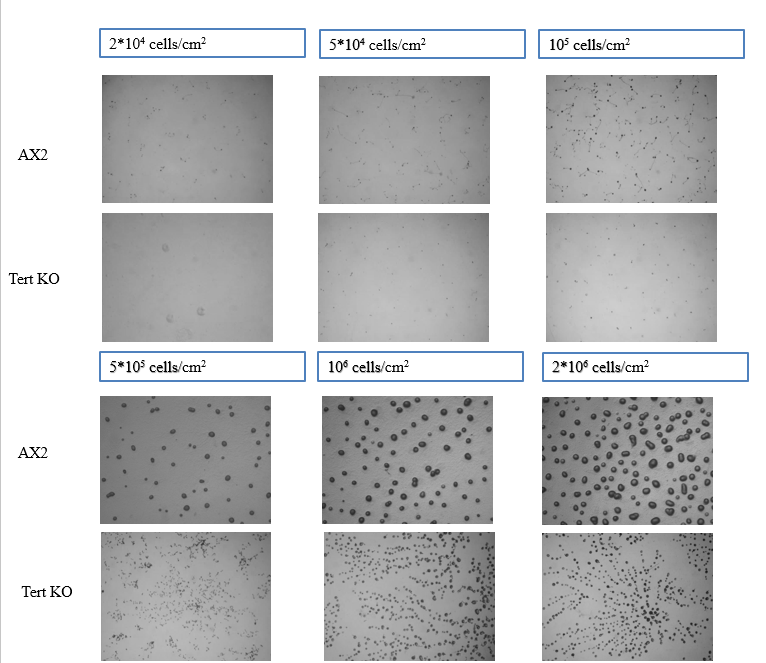

**S3 Fig**

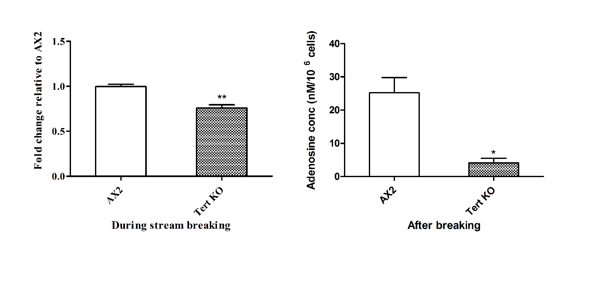

**S4 Fig**

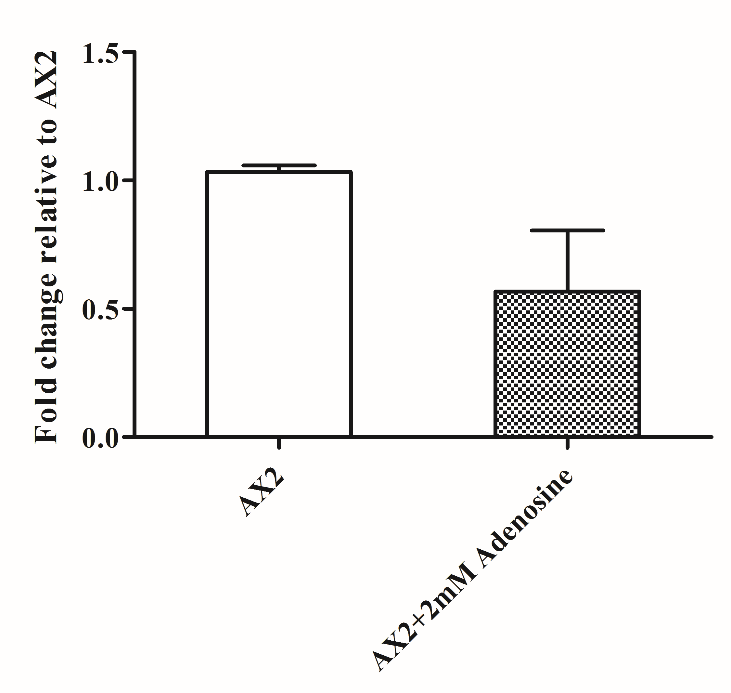

**S5 Fig**

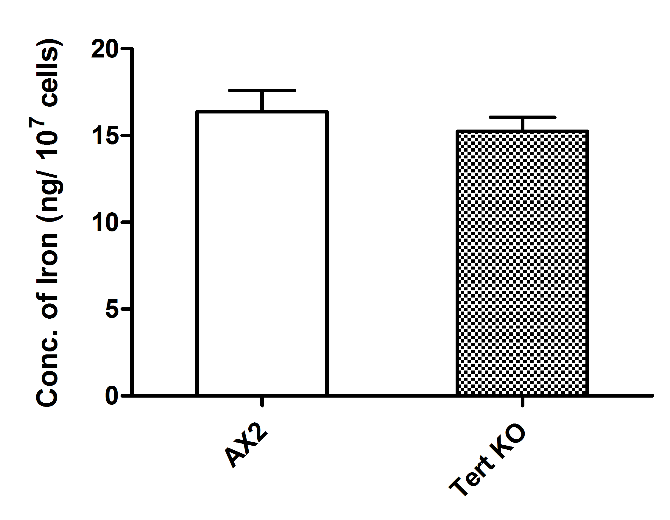

**S6 Fig**

Table S1: Primers used for overexpression vector construction.

| **PRIMER NAME** | **SEQUENCE** |
| --- | --- |
| Tert OE FP | CGGAAGCTTAAAAAAATGAATATTCAAAACATTTATAAAG |
| Tert OE RP | GGCGGTACCATAAATTATATTTAAAATATCAATTG |
| rnlA FP_RT | TCCAAGAGGAAGAGGAGAACTGC |
| rnlA RP_RT | TGGGGAGGTCGTTACACCATTC |

Table S2: Primers used for real-time PCR.

| **PRIMER NAME** | **SEQUENCE** |
| --- | --- |
| tert FP | ACAACAGACAACACTGAAAAG |
| tert RP | CAAAATGTCTTTCTGAAATTC |
| countin FP | CAACCGGTAATGCTTTTGGT |
| countin RP | CACAAACGAGAGCTGACA |
| acaA FP | CATTCTAGAGGCGGTATTGGC |
| acaA RP | GGAGAAAATGTCTGATTTCGCTT |
| carA FP | ATGTTGGGTTGTATGGCAGTG |
| carA RP | AGGGAAACCACCATTGACAG |
| pdsA FP | CCATTGGGTACAACTGGTGGA |
| pdsA RP | AACTGCCCATGATGGATAGGT |
| regA FP | TAAAGCAACGTTGGCACAAG |
| regA RP | ATGGTGATTCCATTGCTTCC |
| pde4 FP | GATCTTGATACACCAATCGAA |
| pde4 RP | CTTCTGCATCATCTGTACATG |
| 5’nt FP | CAGCTGAACAAGTAGCAATGG |
| 5’nt RP | TGGTGGAAGACTTGATGCTG |
| rnlA FP_qRT | TTACATTTATTAGACCCGAAACCAAGCG |
| rnlA RP_qRT | TTCCCTTTAGACCTATGGACCTTAGCG |
